# A *Mecom—Cdk6* toggle switch governs hematopoietic stem-to-multipotent fate transitions via distinct multistable landscapes

**DOI:** 10.64898/2026.06.26.734844

**Authors:** Jonathan A. Martinez, Anupam Dey, MeiLu McDermott, K. Lenhard Rudolph, Adam L. MacLean

## Abstract

Hematopoietic stem cells are required to regenerate the blood system throughout life. We previously discovered that transitions between quiescent stem and multipotent progenitor states are controlled by mutual inhibition between *Mecom* and *Cdk6*, with upstream regulation from the insulin-like growth factor (IGF) signaling pathway. To investigate the dynamics of exit from quiescence and the stem-to-multipotent cell state transition, we modeled the *Mecom*–*Cdk6* regulatory network via coupled nonlinear differential equations. Bifurcation analysis revealed that the model permits tetrastability, with two stable intermediate states, suggesting that stem cell exit from quiescence proceeds via multiple fine-scale transitions. Perturbation of *Mecom* self-activation reorganized the multistable landscape, producing two IGF-dependent landscapes with distinct geometries. At high IGF, transitions proceeded only through an intermediate state, whereas at low IGF, a distinct landscape emerged permitting direct transitions between cell states. Stochastic simulations and minimum action path analysis showed that the multipotent attractor is deep at high IGF, whereas low IGF promotes a transition to quiescence by stabilizing the stem cell state. Simulated pharmacological intervention via CDK4/6 inhibitors destabilized the multipotent state and favored transitions towards a more stem-like quiescent state. Together, these results demonstrate how IGF signaling, *Mecom* self-activation, and *Cdk6* inhibition jointly shape early stem cell fate decisions by dictating the accessible cell states on multistable landscapes and the transition paths that connect them.

## 1 Introduction

Hematopoietic stem cells (HSCs) sustain blood formation by giving rise to a hierarchy of progenitor and differentiated cell types. In the bone marrow, HSCs differentiate into multipotent progenitors (MPPs) and lineage-restricted precursors (MacLean et al., 2017; Mendelson & Frenette, 2014; S. J. Morrison & Scadden, 2014). A defining feature of HSCs is quiescence, a reversible state of cell-cycle exit that preserves stem cell potential (Fujino et al., 2022) and protects against exhaustion (Cheng et al., 2000) or malignant transformation (Wang & Dick, 2005). Transitions between stem and multipotent states must be tightly regulated to maintain long-term function while enabling blood production upon demand. However, the gene regulatory mechanisms that govern these earliest HSC fate decisions remain unclear.

A growing body of evidence identifies Cyclin-dependent kinase 6 (*Cdk6*) and the MDS1 and EVI1 complex locus (*Mecom*) as key regulators of HSC quiescence and activation. *Mecom* promotes HSC quiescence by maintaining stemness, self-renewal, and long-term repopulating capacity (Calvanese et al., 2022; Voit et al., 2023; Yokomizo et al., 2022). In contrast, *Cdk6* promotes exit from quiescence (G0) into the cell cycle by promoting cell-cycle entry and proliferation (Kollmann et al., 2013; Laurenti et al., 2015; Scheicher et al., 2015). Prior computational work has predicted a mutually inhibitory interaction between *Mecom* and *Cdk6*, suggesting that these factors regulate transitions between a *Mecom*-high/*Cdk6*-low stem state and a *Mecom*-low/*Cdk6*-high multipotent state (Rommelfanger et al., 2025). External signals further modulate this balance, including the IGF pathway, which promotes *Cdk6* activity and suppresses *Mecom* expression (MacLean & Rudolph, 2024; Rommelfanger et al., 2025; K. Young et al., 2021; K. A. Young et al., 2024). Together, these findings suggest that *Mecom*, *Cdk6*, and IGF form a regulatory network controlling transitions between stem and multipotent states, but how the structure of this network shapes transition pathways remains unclear.

Early dynamical systems models sought to explain how gene regulatory circuits encode hematopoietic lineage decisions. A classical example is the *GATA1*–*PU.1* toggle switch, in which mutual inhibition and self-activation generate multiple stable states corresponding to progenitor and lineage-committed cell identities (Huang et al., 2007). By analyzing how changes in regulatory interactions alter the underlying attractor landscape through bifurcations, these models demonstrated how cell fate commitment can be driven by the reorganization of a landscape that a bifurcation will induce. Despite a considerable body of literature on such multistable models and the cell fate choices they control, characterization of the earliest cell fate decisions during hematopoiesis as cells exit from a quiescent stem cell state by such means has remained elusive.

Recent studies in ontogeny have adopt a geometric perspective on the gene regulatory control of cell fate landscapes (Raju & Siggia, 2023; Rand et al., 2021; Sáez et al., 2022). Rather than focusing on a specific gene regulatory network and the states that it produces, this framework uses simple gradient-like models and classical tools from dynamical systems theory to examine how perturbations give rise to bifurcations driving cell fate decisions. Bifurcation and catastrophe theory provide the mathematical framework for describing these geometric reorganizations as regulatory parameters vary, giving rise to distinct transition landscapes (Duddu et al., 2025; Raju & Siggia, 2024; Rand et al., 2021). Such approaches have been applied both in the theoretical case and to experimental models, revealing common principles governing cell fate decisions across diverse biological systems (Cislo et al., 2025; A. Howe & Mani, 2025; Sáez et al., 2022; Yampolskaya et al., 2025). Here, we take a geometrical approach to study the *Mecom*–*Cdk6* network and seek to determine how regulatory interactions reorganize transition landscapes and thereby alter early hematopoietic stem cell fate transitions.

To this end, we develop a minimal model of *Mecom*–*Cdk6* mutual inhibition modulated by IGF signaling and analyze its dynamical behavior across signaling and regulatory conditions. We show that the system supports up to four stable states, including intermediate states between stem and multipotent identities, suggesting that transitions between these cell states may proceed through a series of smaller regulatory steps rather than a single commitment event. We further show that weakening *Mecom* self-activation reorganizes the underlying transition landscape, producing distinct dynamical regimes at low and high IGF signaling. While high IGF preserves transitions through intermediate states, low IGF permits qualitatively different transition behaviors that allow more direct switching between cell states. Using two-parameter bifurcation analysis, we then identify broader regions of parameter space associated with distinct transition landscapes and characterize how these landscapes emerge across combinations of IGF signaling and *Mecom* self-activation. Finally, we investigate whether inhibition of *Cdk6* can reshape these transition landscapes, demonstrating that perturbation of the *Mecom*–*Cdk6* network can bias transitions toward stem states.

## 2 Results

### 2.1 A *Mecom*–*Cdk6* toggle switch induces fine-scale transitions through multiple stable intermediate states

To investigate stem cell fate decisions under the control of a *Mecom–Cdk6* inhibitory loop (toggle switch), we developed a differential equation model of *Mecom–Cdk6* dynamics. The regulatory network consists of the core mutual inhibitory motif identified in previous work (Rommelfanger et al., 2025), with upstream regulation from the insulin-like growth factor (IGF) signaling pathway (Fig. 1A). In the network, *Mecom* (*X*) and *Cdk6* (*Y*) both exhibit self-activation. IGF signaling (*S*), represented in the model as a parameter, inhibits *Mecom* and activates *Cdk6*. Our previous work predicted that this network controls early stem cell differentiation from a quiescent hematopoietic stem cell (HSC) state to a multipotent progenitor state, under both normal (ad libitum) and diet-restricted conditions at young age, although the network may be perturbed during hematopoietic aging. The HSC state is characterized by high *Mecom* and low *Cdk6* expression; the multipotent state is characterized by low *Mecom* and high *Cdk6* expression. Here, we seek to understand how these stem cell fate decisions take place under different dietary conditions.

**Figure 1:**
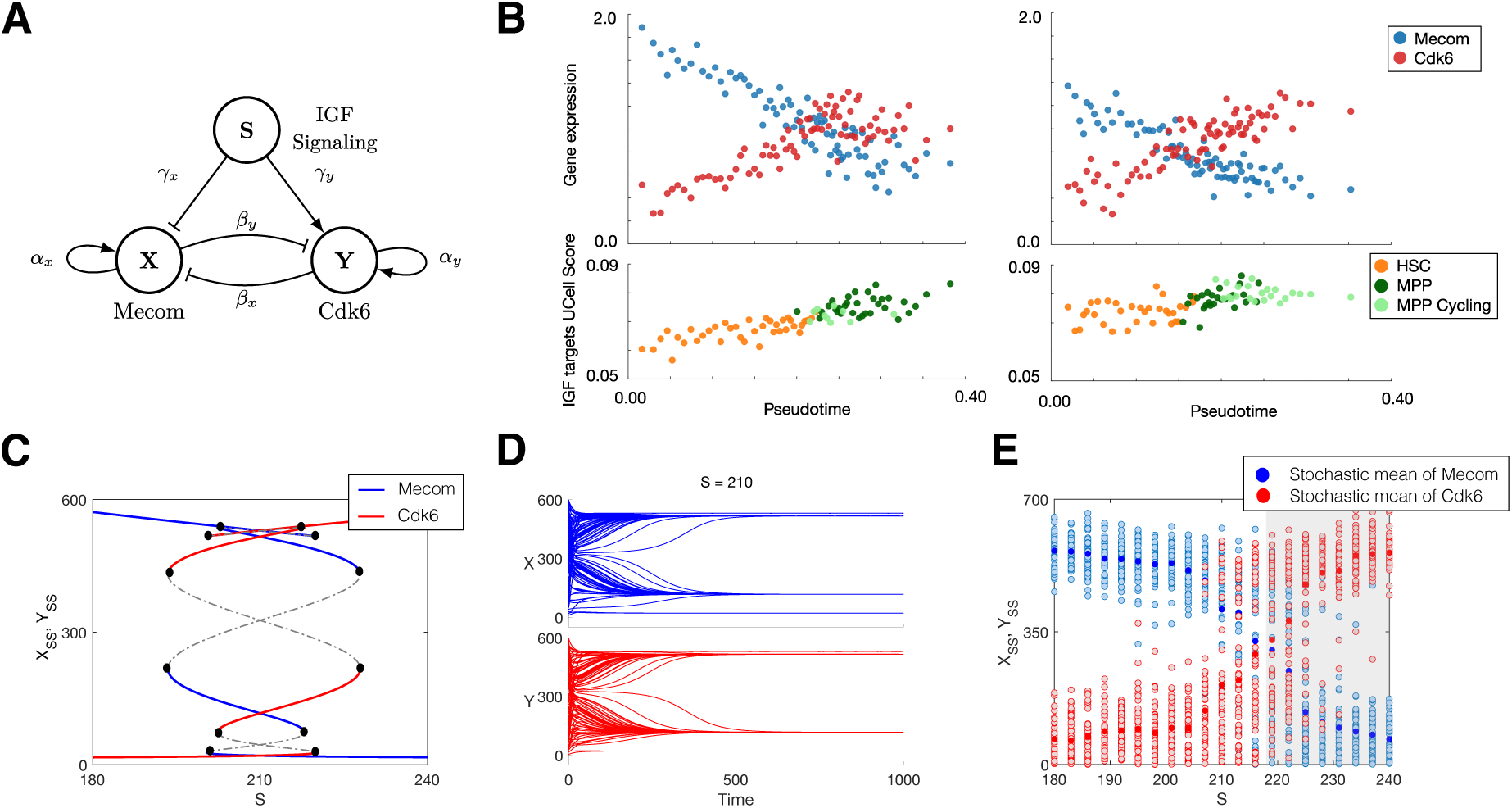
Modeling a Mecom–Cdk6 regulatory network governing stem-to-multipotent transitions. **(A)** Schematic of the regulatory network in which *Mecom* (X) and *Cdk6* (Y) mutually inhibit each other and are regulated by IGF signaling (*S*). **(B)** Single-cell gene expression plotted as pseudocells over pseudotime recapitulates the stem-to-multipotent transition in young mice fed ad libitum (left) or diet restricted (right). Bottom row shows IGF target gene expression as UCell scores over pseudotime, colored by cell type. **(C)** Bifurcation diagram of *Mecom* (X) and *Cdk6* (Y) with respect to *S*. **(D)** Simulation of *Mecom* and *Cdk6* at *S* = 210 from multiple initial conditions. **(E)** Stochastic simulations of the model across varying IGF levels. Light blue/red points denote single realizations from independent simulated trajectories (approximating the stationary distribution). Bright blue/red points denote the mean value of the stationary distribution at each *S*.

We re-analyzed gene expression patterns during the HSC-to-multipotent transition to assess the association between *Mecom*, *Cdk6* and IGF signaling. Whereas previously we studied both cell cycle regressed and cell cycle non-regressed gene expression (Rommelfanger et al., 2025), here we conducted a partial regression to remove only the genes with the strongest cell cycle influence. The hematopoietic cell states identified were the same, albeit with cell cycling distinction in the multipotent pool (Fig. S1). Along pseudotime from stem to committed progenitor states in young age (ad libitum or diet restricted), *Mecom* expression decreased whereas *Cdk6* expression increased (Fig. 1B). In parallel, we observed an increase in IGF signaling pathway target gene expression (Fig. 1B, bottom), as represented by UCell scores for the mTORC1 signaling pathway, which lies downstream of IGF and PI3K/Akt. This association between IGF signaling and *Mecom/Cdk6 expression* across both conditions further motivates the network topology: as IGF signaling increases, cells are driven away from a *Mecom*-high state towards a *Mecom*-low/*Cdk6*-high state.

Bifurcation analysis of the Mecom-Cdk6 model with IGF signaling (*S*) as the bifurcation parameter from an initial parameterization (see Methods) revealed up to four co-occurring stable fixed points, i.e. tetrastability (Fig. 1C). In the tetrastable region, two stable intermediate states exist, situated close to the high and low expression states, respectively (Fig. 1D). Analysis of the basins of attraction on the phase plane showed that the intermediate states separate the high/low expression states and occupy substantially larger basins of attraction than high/low expression states representing stem or multipotent progenitor cells (Fig. S2). The organization of these basins of attraction produces a canalized flow through intermediate regions of the phase space, suggesting that progression from stem cell to multipotent state occurs via transition through stable intermediate states rather than in a single step.

To assess the impact of noisy gene expression dynamics on the cell state transitions, we also performed simulations of the Mecom–Cdk6 network using a stochastic differential equation (SDE) implementation of the model (see Methods). At low IGF signaling (*S*), simulations approximating the stationary distribution converge to *Mecom*-high/*Cdk6*-low states, similarly at high *S*, simulations converge to *Mecom*-low/*Cdk6*-high states with very low probability of state switching (Fig. 1E). However, near the tetrastable region, the stationary distribution for both *Mecom* and *Cdk6* broadens to encompass low, high, and intermediate states. In this region, basins appear to be shallow relative to the noise, permitting frequent transitions between all stable states. Equivalent levels of noise observed experimentally would make it hard or impossible to resolve the multiple stable cell states from the data, rather transitions would appear as a relatively continuous transition through transcriptional space (as is indeed observed). This model in the tetrastable regime makes a specific prediction about hematopoiesis: that HSC-to-multipotent transitions occur not in a single step but rather via multiple fine-scale transitions mediated by stable intermediate states.

### 2.2 Reduced *Mecom* self-activation produces distinct multistable landscapes at low and high IGF

To investigate the impact of *Mecom* self-activation on the multistable HSC-to-multipotent transition landscape, we perturbed the *Mecom* self-activation rate (*α_x_*), which, through the positive feedback loop it forms, promotes high *Mecom* expression and thus stabilizes the stem cell state (Fig. 2A). This perturbation was biologically motivated as a means to study in greater depth the association between *Mecom* self-activation rates and HSC quiescence (Rommelfanger et al., 2025). HSC quiescence is in turn impacted by changes in physiological contexts such as during dietary perturbation or aging (Mejia-Ramirez & Florian, 2020; Tümpel & Rudolph, 2019).

**Figure 2:**
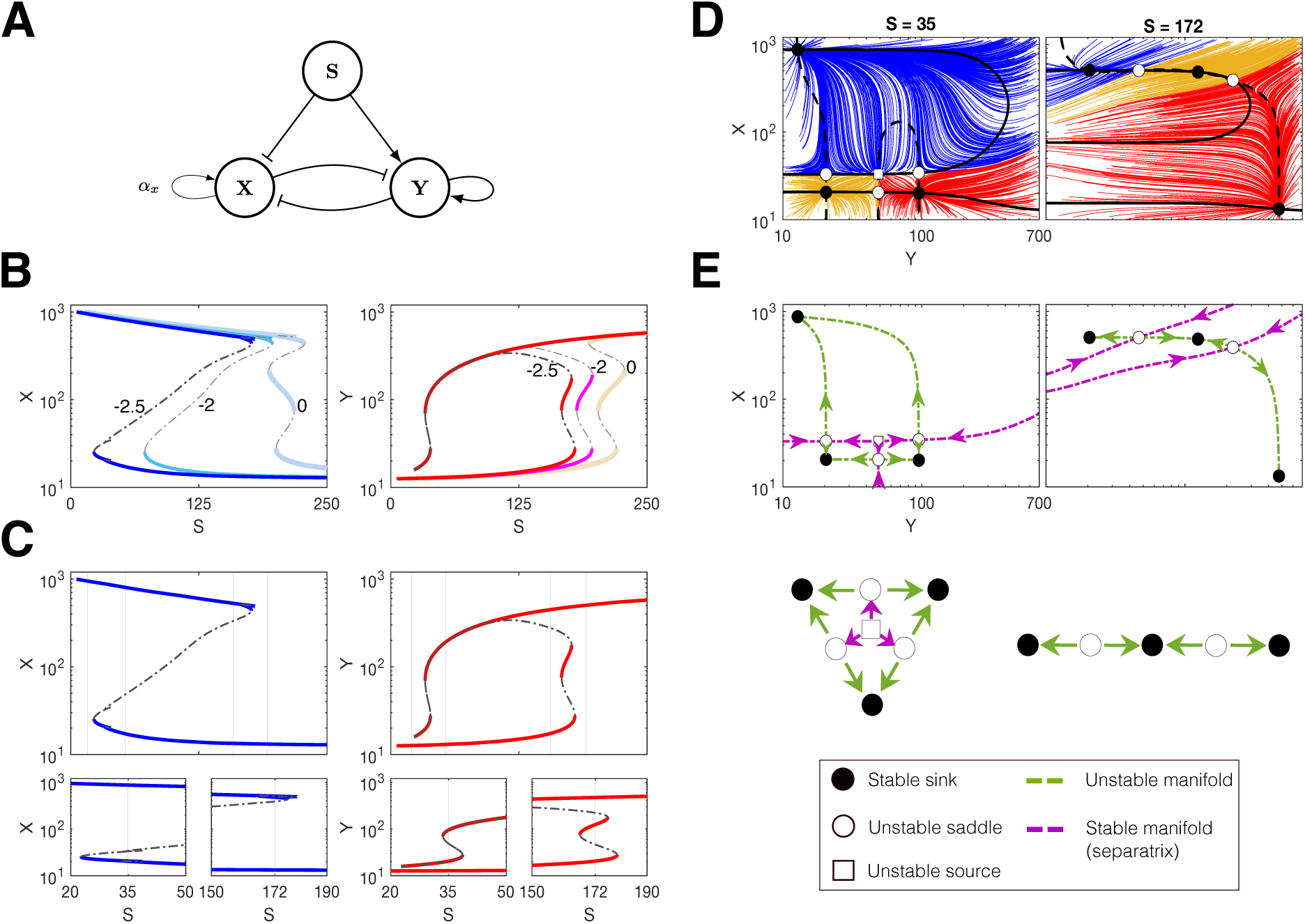
Reduced Mecom self-activation produces distinct multistable landscape. **(A)** Schematic highlighting the reduction in *Mecom* self-activation strength (*α_x_*). **(B)** Bifurcation diagrams of *Mecom* (X) and *Cdk6* (Y) as a function *S* at baseline, 2-fold decrease, and 2.5-fold decrease in *α_x_*. **(C)** Expanded view of the bifurcation plot at 2.5-fold decrease in *α_x_*, insets show zoomed-regions of multistability. **(D)** Phase-plane geometries and trajectory flows at representative IGF values (*S* = 35 and *S* = 172). **(E)** Stable and unstable manifolds defined for the same landscapes as in (D). Bottom: topological arrangement of the saddle-node points of each landscape.

Reducing the *Mecom* self-activation rate reshaped the *Mecom–Cdk6* multistable landscape (Fig. 2B). At baseline, with low IGF signaling only the monostable *Mecom*-high state exists. At a 2-fold reduction in *Mecom* self-activation rate, the model exhibits bistability even at low IGF levels, permitting the coexistence of a *Mecom*-low (*Cdk6*-high) state. Thus, a sufficiently high *Mecom* self-activation rate is required to preserve the exclusive stability of the *Mecom*-high stem cell state.

At a 2.5-fold reduction in *Mecom* self-activation, new stable states emerge, reshaping the landscape more dramatically (Fig. 2C). At high IGF, a new stable intermediate state emerges between *Mecom*-high and *Cdk6*-high states, allowing for fine-scale transitions from stem cell to multipotent states as observed above. At low IGF signaling, a new stable state also emerges, thus creating a new tristable region with a distinct geometry from that observed at high IGF signaling.

The phase planes for these two regions: low (*S* = 35) and high (*S* = 172) IGF signaling are shown in Fig. 2D-E. At low IGF, the *Mecom*-high attractor (blue trajectories in Fig. 2D) occupied the largest region of phase space; only at very low *Mecom* levels do the intermediate and *Cdk6* high states become accessible. Biologically, in this regime, cells are thus highly likely to be locked into a quiescent stem cell state. At high IGF, the *Cdk6*-high attractor (red trajectories) occupies the largest region of phase space, and transitions between the *Mecom*-high and the *Cdk6*-high states proceed via the intermediate state. To highlight the differences in phase planes between these regions and clarify the basin boundaries, we plotted the stable and unstable manifolds of each phase plane (Fig. 2E top). We also sketched the critical points of each phase plane topologically (Fig. 2E bottom). The topology of the fixed points at low IGF resembles the unfolding of an elliptic umbilic bifurcation, one of the catastrophes laid out by René Thom is his seminal theory (Thom, 1975). While it is not strictly an elliptic umbilic, as i) this is not a gradient system, and ii) the bifurcations studied thus far are codim-1; not codim-2 as would be required, the radial symmetry of three stable fixed points around a source represents an elliptic-umbilic-like structure. The relevance of this is that the landscape permits direct transitions between any pair of stable states: characterizing stem cell, multipotent progenitors, and intermediate-state cells.

In contrast, at high IGF, transitions from the *Mecom*-high state to the *Cdk6*-high state occur only via a canalized path passing through the stable intermediate (Fig. 2E). I.e. stem cells exit from quiescence and differentiate into multipotent progenitors indirectly through an intermediate, as was also observed above in the tetrastable regime. Phase planes at multiple IGF signaling levels reveal this landscape is reorganized through several bifurcations as IGF signaling increases from low to high values (Fig. S3).

Two additional parameter sets that each permitted tetrastability were studied in order to assess the generality of the results observed. For both these additional parameter sets, reducing the strength of the *Mecom* self-activation led to the same qualitative bifurcations whereby distinct multistable landscapes emerged at low and high IGF signaling (Fig. S4). Overall, we have found that the quiescent stem cell state is sensitive to *Mecom* self-activation: reducing this parameter does lead to the emergence of a *Cdk6*-high state even at low IGF. However, the quiescent state remains highly stable at low IGF. In contrast, even with reduced *Mecom* self-activation, at high IGF transitions from stem to multipotent states that are canalized through a stable intermediate are preserved.

### 2.3 Transition paths reveal the IGF-dependence of deep stem cell attractors that are difficult to escape

To investigate the dynamics and transition paths across the landscapes identified above (Fig. 2D), we analyzed stochastic simulations of the *Mecom-Cdk6* network initialized from different basins. At high (*S* = 172) IGF signaling levels (Fig. 3A–C), the *Mecom*-high attractor was found to be relatively shallow. Trajectories initialized at the *Mecom*-high state transitioned out of this state: at low noise remaining long-term in the intermediate state (Fig. 3A) and at high noise remaining in the *Cdk6*-high state (Fig. 3B). Trajectories initialized at the *Cdk6*-high state remained in this state long-term regardless of noise level. These simulations reveal the relatively shallow depth of the *Mecom*-high attractor state at high IGF signaling: cells will transition out of a stem cell state into a multipotent state readily here. Qualitatively similar behaviors were observed across a broad range of noise strengths studied, with trajectories progressively accumulating in the intermediate and *Cdk6*-high states as noise increased (Fig. S5A–D). We also analyzed the most probable transition paths via the Freidlin-Wentzell functional and associated minimum action paths (MAPs). We found that transitions out of the *Mecom*-high state proceed through the intermediate state, i.e. indicating that cells follow canalized paths on this landscape through the stable intermediate state en route to the *Cdk6*-high multipotent state (Fig. 3C).

**Figure 3:**
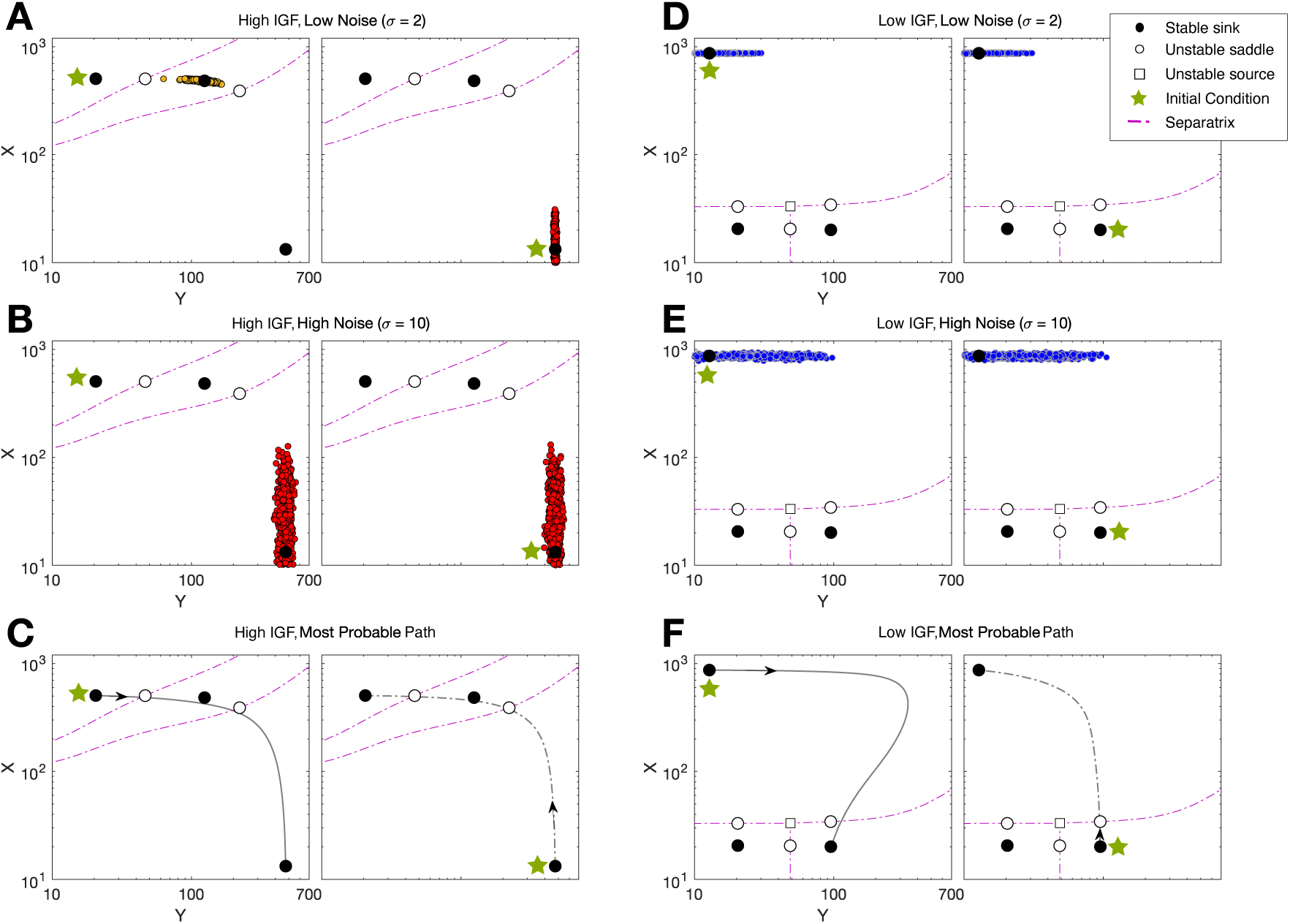
Noise-driven transitions at high and low IGF regimes. **(A–B)** High IGF signaling regime (*S* = 172). Stochastic trajectories are initialized at the *Mecom*-high (left) or *Cdk6*-high (right) states under low noise (*σ* = 2, A) or high noise (*σ* = 10, B). Colored points show the cell states at the final simulation time (1000 cells at time *t* = 10^4^). **(C)** Minimum action paths for *S* = 172, showing transitions from the *Mecom*-high state to the *Cdk6*-high state (left) and from the *Cdk6*-high state to the *Mecom*-high state (right). **(D–E)** Low IGF signaling regime (*S* = 35); as for (A–B) with low noise (D) and high noise (E). **(F)** Minimum action paths for *S* = 35, shown as in (C).

In contrast, at low (*S* = 35) IGF signaling (Fig. 3D–F), the relative stability of the *Mecom*-high and *Cdk6*-high basins of attraction is reversed. At low IGF, trajectories initialized near the *Mecom*-high state remain confined to this state under any noise level simulated. Trajectories initialized near the *Cdk6*-high state transition out of this state and immediately into the *Mecom*-high state even at low noise levels. This preference for the *Mecom*-high state persisted across all simulated noise strengths, with the distribution of cell states overwhelmingly concentrated in the stem-state basin despite substantial stochastic fluctuations (Fig. S5E–H). MAPs calculated for this landscape show that transitions into/out of the *Mecom*-high state proceed directly, without passing through the intermediate state, as permitted by the landscape geometry (Fig. 2D-E). Thus, as low IGF signaling levels cells become locked into a *Mecom*-high, stem cell state, requiring very high noise or external perturbation to exit from quiescence and differentiate.

IGF signaling thus creates a directional fate bias. High IGF favors the *Cdk6*-high multipotent progenitor cell state, whereas low IGF favors the *Mecom*-high stem cell state. The high IGF condition modeled here likely corresponds to homeostatic hematopoiesis, i.e. mice fed ad libitum diet in (Rommelfanger et al., 2025). Under these conditions the majority of hematopoietic production proceeds via multipotent progenitors. Low IGF then may correspond to the conditions observed with diet restriction (which reduces metabolic activity and therefore IGF levels) whereby propensity for stem cell quiescence increases.

### 2.4 Two-parameter bifurcation analysis reveals elliptic umbilic-like geometries only at low IGF

So far, we have examined how IGF signaling (*S*) and *Mecom* self-activation (*α_X_*) shape the *Mecom*–*Cdk6* landscape at selected parameter values (Figs. 2-3). To extend this analysis across a broader parameter space, we studied two-parameter bifurcations across the (*S, α_X_*) plane (Fig. 4A). For intermediate-high levels of IGF signaling, i.e. *S* & 100, we observe many different multistable regions as the several saddle-node curves are crossed, as well as co-dim 2 cusp points where these curves collided, which we confirm via properties of the quadratic and cubic coefficients (see Methods). At low IGF, over a range of *α_X_*values, a source node appears, and with the emergence of an elliptic umbilic-like bifurcation within saddle-source region (Fig. 4A inset, and Fig. S6A-B).

**Figure 4:**
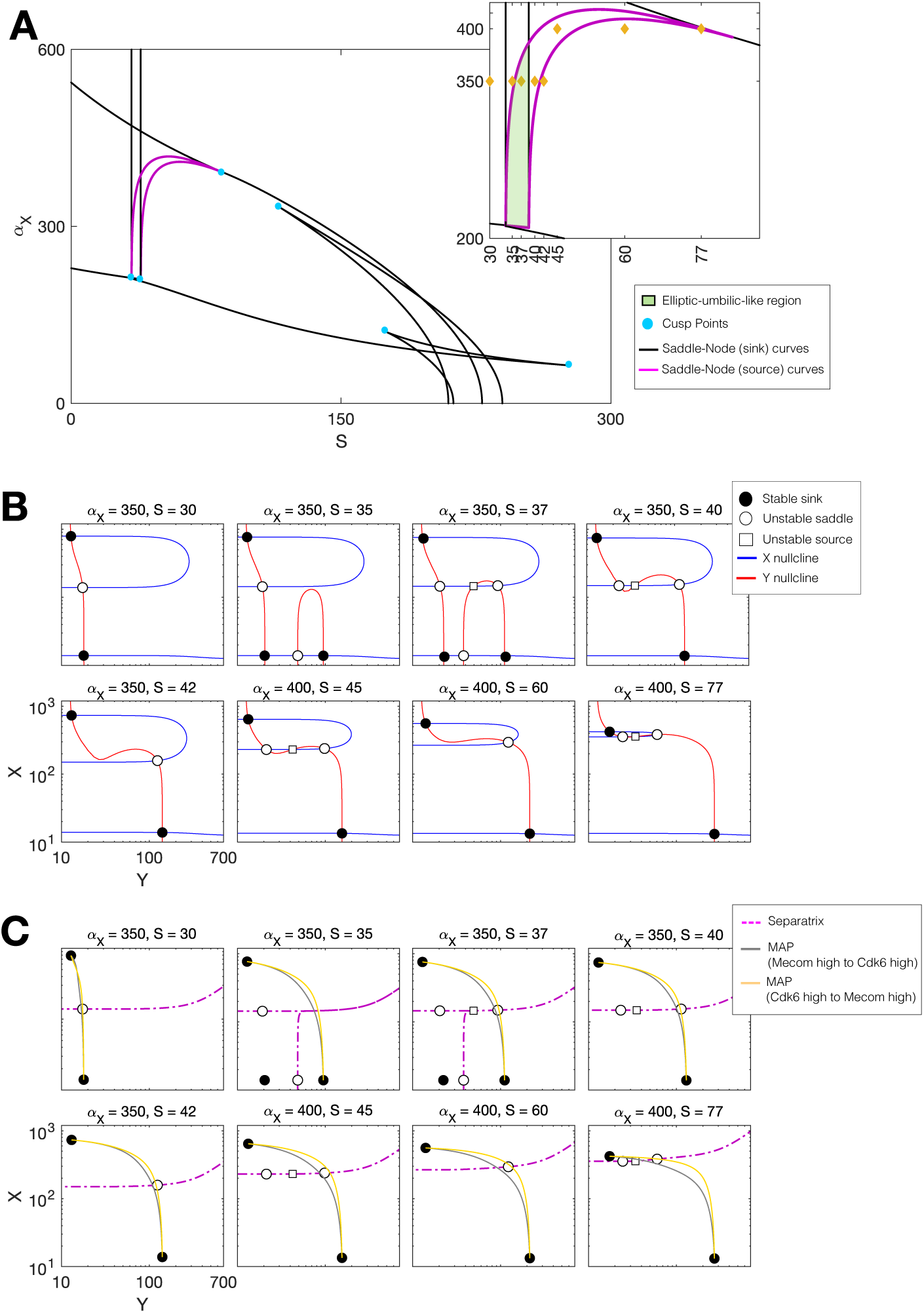
Two-parameter bifurcation analysis reveals distinct landscapes and the transitions they permit. **(A)** Two-parameter bifurcation diagram in (*S, α_X_*). Inset: zoomed region, showing parameter sets plotted below (gold diamonds). **(B)** Phase planes at representative parameter values illustrating the configuration of fixed points at each. **(C)** Most probable paths corresponding to the parameter sets in (B), depicting transitions between Mecom and Cdk6 high steady states.

To characterize how these different landscapes influence cell state transitions in the low IGF regime, we examined representative paths through landscape space at *α_X_* = [350, 400], together with the corresponding most probable paths on each landscape (Fig. 4B-C). At *α_X_*= 350 and very low IGF (*S* = 30), the landscape consists of only stem and intermediate states. As IGF increases to *S* = 35, the multipotent state emerges, requiring cells to transition through the stable intermediate state before reaching the multipotent state and producing the canalized transition geometry described in Sections 2.2–2.3. A further increase in IGF (*S* = 37) reorganizes the landscape into an elliptic-umbilic-like geometry, allowing cells to transition directly between all three stable states. At still higher IGF signaling (*S* = 40-42), the intermediate state and subsequently the source are lost through successive bifurcations, leading to a bistable landscape in which only the stem and multipotent states remain. At *α_x_* = 400 for higher IGF signalling (*S* = 45-77), we see this source reappear before reaching a cusp. One-parameter bifurcation plots (cross sections of the two-parameter bifurcation) show how the source is created and destroyed (Fig. S6C–D).

Together, these results show that direct transitions between stem and multipotent states are restricted to low IGF signaling, whereas higher IGF signaling preserves canalized differentiation through stable intermediate states.

### 2.5 Pharmacological intervention via simulated inhibition of *Cdk6* destabilizes the multipotent state

Given the importance of *Cdk6* inhibition in cancer therapy (CDK4/6 inhibitors are part of standard-of-care combination therapy for certain breast cancer subtypes) (L. Morrison et al., 2024), we simulated the effects of *Cdk6* inhibition to assess the impact on the stem cell— multipotent landscapes. To mimic the effects of CDK4/6 inhibitors (e.g. via Palbociclib, Ribociclib, or Abemaciclib (Braal et al., 2021), although we remain drug agnostic here), we performed stochastic simulations of cells initialized at the *Cdk6*-high state at high (*S* = 172) IGF signaling, with increasing values of the *Cdk6* degradation rate representing increasing drug doses (Fig. 5A). In the absence of drug (*d_drug_*= 0), the landscape is tristable and cells initialized at the *Cdk6*-high state remain confined there, indicating that the multipotent state is highly stable under these conditions. At *d_drug_*= 0.03, the landscape changes, eliminating the intermediate state. Over long times we see co-occupancy of cells in both *Mecom*-high and *Cdk6*-high states, i.e. enabling a fraction of cells to transition readily between multipotent and stem cell states. As CDK4/6 inhibition increases further (*d_drug_* = 0.06), we no longer see co-occupancy. I.e. over long times all cells will transition out of the multipotent state (the multipotent attractor has become sufficiently shallow, relative to the noise level). At *d_drug_*∈ [0.09, 0.15], the landscape changes once again: we see the emergence of a stable intermediate state with an elliptic umbilic-like geometry, although neither the intermediate nor *Cdk6*-high basins are deep enough relative to the *Mecom*-high state to retain cells long-term. Finally, at the highest dose considered (*d_drug_* ≥ 0.17), the *Cdk6*-high attractor is destroyed, leaving only stem and intermediate states. These results highlight how the Mecom–Cdk6 toggle switch is druggable. CDK4/6 inhibition destabilizes the *Cdk6*-high multipotent state and deepens the stem cell attractor, eventually eliminating the multipotent state entirely, thus locking cells into a quiescent stem cell state.

**Figure 5:**
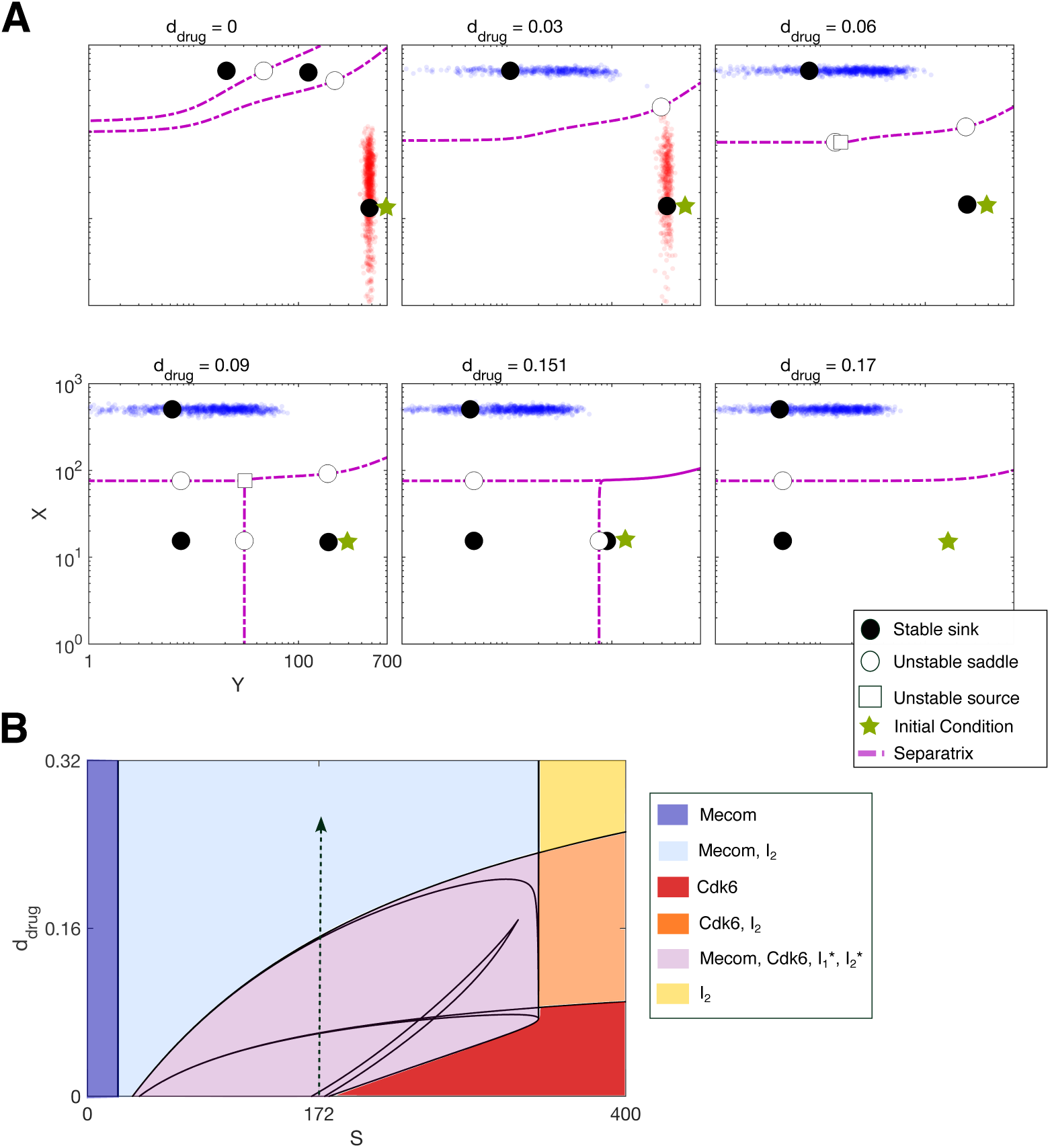
Palbociclib reorganizes the Mecom–Cdk6 landscape to promote transitions toward Mecom-high states. **(A)** Stochastic trajectories on phase planes corresponding to increasing CDK4/6 inhibition (*d_drug_*) at fixed IGF signaling (*S* = 172), initialized at the *Cdk6*-high attractor (green star). Each panel shows 1000 stochastic trajectories evaluated at the final simulation time (*t* = 10^4^). **(B)** Two-parameter bifurcation diagram in (*S*,*d_drug_*) for 2.5-fold reduction in *Mecom* self-activation. Colors denote selected regions of specific steady state configurations. Dashed arrow denotes the path corresponding to the increasing drug concentrations shown in (A).

To assess the impact of CDK4/6 inhibition over a broader range of IGF signaling, we extended the virtual pharmacological analysis to include a range of IGF signaling values and drug doses on a two-parameter bifurcation plot (Fig. 5B), which is colored by the type of stable states that exist in each region. The full set of phase planes used to define the states & region colors are given in Fig. S7A-B. At sufficiently low IGF signaling, even in the absence of drug, no *Cdk6*-high state exists. At intermediate levels of IGF signaling, stem cell and multipotent cell states coexist along with various intermediates, allowing CDK4/6 inhibition to progressively destabilize the multipotent state and promote transitions toward a stem cell state. At high-to-ultrahigh IGF signaling, the multipotent state persists even with the addition of large drug doses; high IGF signaling buffers the multipotent state against pharmacologic CDK4/6 inhibition. Similar transition landscapes are observed in the baseline model with stronger *Mecom* self-activation (Fig. S7C-D). Thus, the ability of CDK4/6 inhibition to destabilize the multipotent state depends on the surrounding IGF signaling environment. At the highest IGF signaling levels and drug doses, we see that both the stem cell and the multipotent stable states are destroyed, leaving only one stable intermediate. Thus we can quantify the limits of the system: parameters values in this region likely represent states outside of the physiological ranges necessary to maintain hematopoiesis.

## 3 Discussion

Here we have developed a mathematical model to investigate how a *Mecom—Cdk6* regulatory network controls transitions from quiescent stem cell to multipotent progenitor cell states. We found that transitions from stem to multipotent states under the *Mecom—Cdk6* toggle switch proceed via fine-scale transitions through one or more stable intermediate states, are shaped by the IGF signaling environment, and are druggable through *Cdk6* inhibition. These results both support and extend our fundamental understanding of early HSC fate decisions. In the classical view of hematopoiesis, differentiation proceeds from long-term to short-term HSCs, followed by subsequent transition to multipotent and then lineage-restricted progenitors (Yang et al., 2005). Prior to the advent of high-throughput single-cell technologies, experimental studies had identified several discrete HSC subsets, including dormant, non-dormant, and intermediate-term stem cell populations (Benveniste et al., 2010; Wilson et al., 2008). The advent of single-cell genomics has since revealed substantially greater heterogeneity within the hematopoietic stem and progenitor cell compartment, fueling debate over whether early hematopoietic differentiation proceeds continuously or through discrete intermediate states (MacLean et al., 2018; Moris et al., 2016; Safina & van Galen, 2024; Sparta et al., 2022). Recent single-cell studies support discrete differentiation suggest that several intermediate states are present along the HSC-to-MPP transition (Ediriwickrema et al., 2025; Jakobsen et al., 2024; Weng et al., 2024; Zeng et al., 2026). Our *Mecom—Cdk6* toggle-switch model provides one possible mechanistic explanation for these observations by showing how interactions between *Mecom*, *Cdk6*, and IGF signaling can give rise to intermediate states during HSC-to-MPP differentiation.

The *Mecom—Cdk6* toggle-switch model suggests that HSC quiescence is regulated by the balance between *Mecom* and *Cdk6* activity, generating two experimentally testable hypotheses concerning how the effects of aging and metabolism are impacted by *Mecom* and *Cdk6*. *Mecom* has been implicated in the maintenance of HSC quiescence, a defining feature of long-term stem cells that is closely linked to self-renewal capacity (Nakamura-Ishizu et al., 2014; Rommelfanger et al., 2025). Maintenance of HSC quiescence becomes progressively impaired during aging, contributing to a decline in stem cell self-renewal capacity (de Haan & Lazare, 2018; Mejia-Ramirez & Florian, 2020; Tümpel & Rudolph, 2019). Our model predicts that aging-associated loss of quiescence may arise through weakening *Mecom* self-activation. In turn, weakened *Mecom* self-activation will permit quiescent HSCs to differentiate under metabolic conditions that would otherwise maintain stem cell quiescence. This hypothesis could be tested *in vivo* by perturbing *Mecom* transcriptional activity and assessing HSC quiescence, and aging-associated phenotypes such as myeloid skewing and transplantation capacity. In contrast, *Cdk6* promotes exit from quiescence (Maurer et al., 2021; Scheicher et al., 2015). Recent genetic studies have demonstrated that CDK6 inhibits HSC quiescence in a kinase-independent manner (Mayer et al., 2024). Pharmacological inhibition of CDK4/6 has also been shown to preserve HSC function during hematopoietic stress by increasing HSC quiescence prior to chemotherapy (He et al., 2017; Johnson et al., 2010). These observations motivated our analysis of CDK4/6 inhibition in the *Mecom—Cdk6* toggle-switch. The model predicted that as CDK4/6 inhibiton increased, the ability to differentiate was progressively lost until cells become locked into a quiescent stem cell state. This hypothesis could be tested experimentally by treating HSPCs with CDK4/6 inhibitors and measuring, as above, markers of HSC quiescence and aging-associated phenotypes.

Some limitations of this study highlight important future directions for work. Given the idealized formulation of the model — consisting of a tightly constrained number of transcriptional parameters — pharmacological interventions and the action of IGF signaling on gene expression are represented as effective parameters, rather than fully described. Consequently, only qualitative inferences can be made. In future work, models could incorporate translational and post-translational dynamics to better characterize signal transduction pathways and the cell cycle. However, to expand the model as such successfully will likely require richer quantitative data e.g. from spatial transcriptomics or proteomics. The present model does not explicitly consider aging and its impact on the regulatory network, which was well-supported by the data on at young age (Rommelfanger et al., 2025). Phenomenologically, weakening the *Mecom* self-activation rate was used to represent the potential loss of quiescence capacity with age. In future work, experiments targeting age-associated regulators such as *Igf2bp2* (Suo et al., 2022), together with perturbation of epigenetic regulators (such as DNMT3A and TET2) that are implicated in hematopoietic aging (Kramer & Challen, 2017; Kreger et al., 2024), could help to determine whether aging primarily alters the strengths of existing regulatory interactions or instead rewires the underlying gene regulatory network. HSCs do make fate decisions alone, but rather in the highly specific microenvironments of the bone marrow niche (Grockowiak et al., 2023; MacLean et al., 2017; Rommelfanger & MacLean, 2021; Toghani et al., 2025). This work considers only autonomous fate decisions and future extensions ought to incorporate the impact of niche signaling into regulatory networks.

Taken together, this work establishes a framework for understanding how the *Mecom—Cdk6* toggle switch network governs transitions between quiescent stem cell and multipotent progenitor cell states. Exit from the quiescent stem cell state is controlled by *Mecom* self-activation together with IGF signaling. The network motif (mutual inhibition plus self-activation) may extend beyond *Mecom* and *Cdk6* to provide means for biological networks to build multistable landscapes with key states near the bifurcation point, such that shifts in physiological parameters induce changes to the landscape and thus the cell states transition paths.

## 4 Material and Methods

### 4.1 Mathematical modeling of *Mecom—Cdk6* toggle switch

We modeled the dynamics of *Mecom* (*X*) and *Cdk6* (*Y*) using a system of ordinary differential equations describing the production and degradation of each transcriptional component. Production terms were formulated as Hill-type functions of upstream regulators and IGF signaling, with degradation modeled as linear first-order decay. Variables *X* and *Y* represent effective expression levels.

The model is described be the following equations:

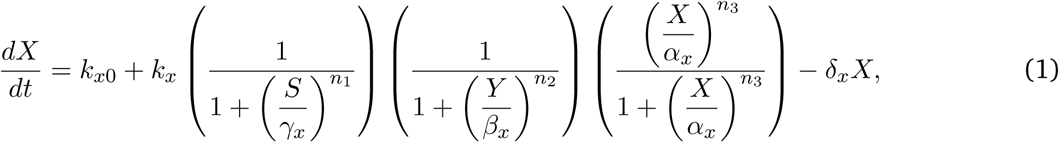

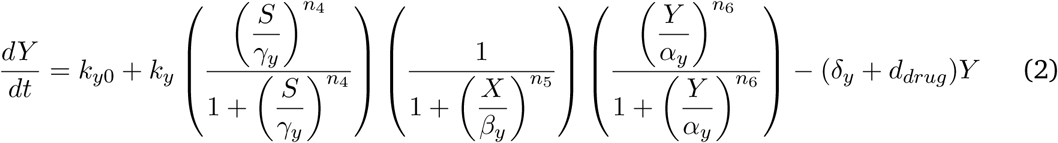

where *k_x_*_0_ and *k_y_*_0_ represent the basal expression of *Mecom* and *Cdk6* in the absence of regulatory inputs; *k_x_* and *k_y_* define the maximal regulated production capacity under fully permissive conditions (see Table 1 for full description). Linear degradation rates *δ_x_* and *δ_y_* capture turnover of *Mecom* and *Cdk6*, respectively. Regulatory interactions are controlled by threshold parameters *α_x_* and *α_y_* governing self-activation, *β_x_*and *β_y_*for the strength of mutual inhibition between *Mecom* and *Cdk6*, and IGF signaling thresholds *γ_x_*and *γ_y_*for the sensitivity of each gene to external signaling input *S*. Hill coefficients *n*_1_–*n*_6_ define the steepness of each regulatory interaction.

**Table 1:** Description of model parameters and baseline values.

| Notation | Description | Baseline Value |
| --- | --- | --- |
| $k_{x0}$ | basal production rate of *Mecom* | 1 |
| $k_{y0}$ | basal production rate of *Cdk6* | 1 |
| $k_x$ | maximal regulated production rate of *Mecom* | 85 |
| $k_y$ | maximal regulated production rate of *Cdk6* | 85 |
| $\alpha_x$ | threshold for *Mecom* self-activation | 87 |
| $\alpha_y$ | threshold for *Cdk6* self-activation | 87 |
| $\beta_x$ | inhibition threshold of *Cdk6* acting on *Mecom* | 392 |
| $\beta_y$ | inhibition threshold of *Mecom* acting on *Cdk6* | 392 |
| $\gamma_x$ | IGF inhibition threshold acting on *Mecom* | 210 |
| $\gamma_y$ | IGF activation threshold acting on *Cdk6* | 210 |
| $n_1$ | Hill coefficient (IGF → *Mecom* inhibition) | 1 |
| $n_2$ | Hill coefficient (*Cdk6* → *Mecom* inhibition) | 3 |
| $n_3$ | Hill coefficient (*Mecom* self-activation) | 2 |
| $n_4$ | Hill coefficient (IGF → *Cdk6* activation) | 1 |
| $n_5$ | Hill coefficient (*Mecom* → *Cdk6* inhibition) | 3 |
| $n_6$ | Hill coefficient (*Cdk6* self-activation) | 2 |
| $\delta_x$ | degradation rate of *Mecom* | 0.08 |
| $\delta_y$ | degradation rate of *Cdk6* | 0.08 |
| $d_{drug}$ | Palbociclib-induced degradation rate of *Cdk6* | 0 |

To model inhibition of CDK4/6 (e.g. via palbociclib or similar drugs), the parameter *d*_drug_ represents the effective increase in *Cdk6* degradation induced treatment. In the absence of drug, *d*_drug_ = 0; positive values correspond to doses of drug that effectively increase the degradation of *Cdk6*.

Regulatory inputs were combined multiplicatively (with AND logic (Dey & MacLean, 2025)), such that high gene expression requires the combination of sufficient self-activation, relief from mutual inhibition, and appropriate IGF signaling.

### 4.2 Bifurcation analysis and phase plane construction

Bifurcation and phase-plane analyses were performed to characterize steady states and their stability as model parameters were varied. Writing the model generically as a two-dimensional system in (*X, Y*) then we have:

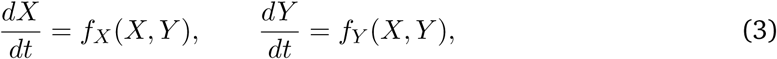

where a fixed point (*X*^∗^*, Y* ^∗^) satisfies the conditions

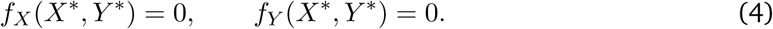

Fixed points were estimated numerically in Matlab (MathWorks) using the function fsolve, which employs a Newton–type root–finding method to solve the coupled nonlinear equations simultaneously. This procedure requires an initial guess (*X*_0_*, Y*_0_), from which successive corrections are computed and used to update the solution iteratively. Iterations were terminated when the norm of the correction satisfied a prescribed tolerance (e.g., ‖ Δ‖ *< ε*). Multiple initial guesses were used to ensure identification of all relevant equilibria within a given parameter regime.

Nullclines were computed by evaluating *f_X_*(*X, Y*) and *f_Y_* (*X, Y*) on a dense two–dimensional grid spanning the state space. The *X*–nullcline was defined by the set of points satisfying *f_X_*(*X, Y*) = 0, and the *Y* –nullcline by *f_Y_* (*X, Y*) = 0. Zero–level contours of these functions were extracted and visualized using the Matlab function contour by specifying the contour level 0. Intersections of the nullclines correspond to fixed points of the dynamical system and provide a direct visualization of equilibrium structure in the phase plane.

Local stability was assessed by linearizing the system about each equilibrium. The Jacobian matrix

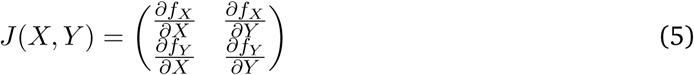

was evaluated at each fixed point using the function Jfun, which returns the Jacobian at a specified state. Eigenvalues of the Jacobian were computed using Matlab’s eig function. Equilibria were classified as stable if all eigenvalues had negative real parts, unstable if all eigenvalues had positive real parts, and saddle points if eigenvalues had mixed signs. These classifications were verified by direct numerical integration of trajectories initialized in the vicinity of each equilibrium.

Stable and unstable manifolds of saddle points were computed by linearizing the dynamical system about each equilibrium and integrating trajectories along the corresponding eigendirections. For each saddle point, eigenvectors associated with the most negative real eigenvalue (dominant stable direction) and the most positive real eigenvalue (dominant unstable direction) were identified. Small perturbations were applied along the positive and negative directions of these eigenvectors to generate initial conditions near the equilibrium. Trajectories initiated along stable directions were integrated backward in time to trace stable manifolds, whereas trajectories initiated along unstable directions were integrated forward in time to trace unstable manifolds. Numerical integration for manifold tracing was performed in Matlab via ode45. The resulting invariant manifolds provide a global representation of separatrices that delineate basin boundaries and organize the phase–space dynamics.

Bifurcation diagrams were computed using the AUTO continuation engine within XPPAUT, which employs pseudo-arclength continuation to trace branches of equilibria as a function of the chosen parameter, starting from a known fixed point at an initial parameter value. For two-parameter bifurcation analysis, a one-parameter continuation was first performed with respect to one parameter to identify saddle-node (fold) points along the equilibrium branch. Each detected fold point was then used as a starting solution for two-parameter continuation, tracing the locus of fold bifurcations as a curve in the two-parameter plane. The tangential meeting point of two such fold curves in the two-parameter plane identifies the cusp bifurcation point.

For the generic two-dimensional system described in Equation 3, with (*a, b*) as two control parameters, a cusp bifurcation point (*X*^∗^*, Y* ^∗^*, a*^∗^*, b*^∗^) was identified by verifying four standard conditions (Kuznetsov, 2023). First, the candidate point must be an equilibrium of the system, meaning that both governing equations vanish at that point. Second, the Jacobian matrix evaluated at the equilibrium must possess a single zero eigenvalue, indicating that the system lies at the boundary between different stability regimes. Third, the leading quadratic nonlinearity along the corresponding critical direction must vanish, distinguishing a cusp from an ordinary fold (saddle-node) bifurcation. Finally, the leading cubic term must remain nonzero, ensuring that the degeneracy is of exactly codimension two and giving rise to the characteristic cusp geometry in parameter space. Together, these conditions provide a rigorous numerical criterion for identifying cusp bifurcation points and locating the intersections of fold branches in the two-parameter bifurcation diagram.

### 4.3 Stochastic simulation via a stochastic differential equation model and minimum–action paths

To investigate the stochastic dynamics of the *Mecom—Cdk6* network model, the deterministic system was extended to a stochastic differential equation (SDE) model of the form

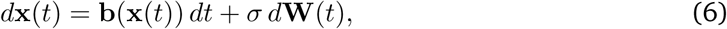

where **x**(*t*) = (*X*(*t*)*, Y* (*t*))^T^, **b**(**x**) = (*f_X_*(*X, Y*)*, f_Y_* (*X, Y*))^T^ is defined by Eqs. (1-2), and **W**(*t*) is a two-dimensional Wiener process with independent components. *σ* represents the noise strength. The SDE model was simulated numerically using the Euler-Maruyama scheme, where we assumed an Itô interpretation.

Trajectories initiated near a stable fixed point typically remained close to that attractor, with occasional noise–driven excursions and rare transitions to other basins of attraction. In the small–noise limit (*σ* → 0), such rare transitions concentrate around a most probable path. The most probable transition path between two attractors was computed by minimizing the Freidlin-Wentzell action functional

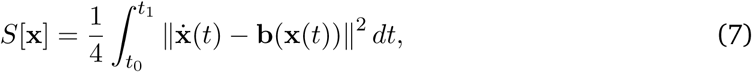

which quantifies the likelihood of a trajectory connecting the initial and final states. In practice, the path was discretized into a sequence of nodes 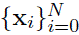 connecting the initial and terminal attractors. The time derivative was approximated using finite differences,

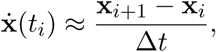

yielding a discretized action functional

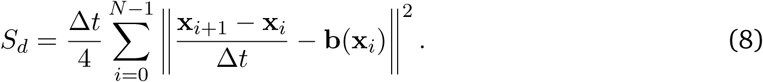

In our implementation, the discretized action was evaluated using the Matlab function computeAction_noD, which computes the numerical approximation of the Freidlin–Wentzell action along a discretized path by penalizing deviations between the path velocity and the deterministic drift. The resulting nonlinear optimization problem was solved using the constrained optimizer fmincon, which minimized the discretized action with respect to the interior path points while keeping the endpoints fixed. The resulting minimum–action path identifies the dominant noise–induced transition route between attractor states, typically passing near saddle points that act as effective transition barriers in phase space.

Minimum–action paths were overlaid with ensembles of stochastic trajectories to compare predicted optimal transition routes with simulated noise–driven dynamics.

### 4.4 Single-cell RNA sequencing data analysis

Single-cell multiomic data (paired snRNA-seq and snATAC-seq; GEO: GSE229892) were obtained from a publicly-available dataset previously generated by our group (Rommelfanger et al., 2025). Bone marrow was collected from 6-month-old female C57BL/6J mice maintained under *ad libitum* feeding or one week of dietary restriction (30% caloric reduction), respectively termed yAL and yDR. LSK cells (Lin^—^Sca-1^+^c-Kit^+^) were FACS-sorted following c-Kit enrichment by magnetic bead selection, and nuclei were isolated for joint single-nucleus multiome sequencing (10x Genomics Chromium). Raw sequencing reads were aligned to the GRCm38 reference genome using Cell Ranger ARC (v2.0.0; 10x Genomics).

scRNA-seq data were processed and analyzed using Scanpy (Wolf et al., 2018). Cells were filtered by a UMI vs ranked barcode metric (yAL: 10 to 4000 barcodes, yDR: 5 to 3600 barcodes), resulting in a total counts distribution from 972-20651 and 1061-11742 respectively. Filters were applied with a minimum of 200 genes per cell and 3 cells per gene. Outliers were identified using median absolute deviation (MAD) of 5 for total counts, number of genes, percent counts in top 50 genes, percent mitochondrial, and percent ribosomal genes. Mitochondrial genes were removed. Scrublet was performed (Wolock et al., 2019). Counts were normalized to 10, 000 and *log*(*x* + 1) transformed. Cell cycle was scored using scanpy’s score_genes_cell_cycle, based on the method developed by Seurat that uses genes from (Satija et al., 2015; Tirosh et al., 2016). Highly variable genes were selected and scaled. Cell cycle genes were then removed from highly variable genes using Mouse GOBP_CELL_CYCLE list of genes from MSigDB (Liberzon et al., 2015). Principal component analysis (PCA) was performed; the top 15 components were used to construct a nearest neighbor graph. Cells were visualized in two dimensions using UMAP (McInnes et al., 2018). Silhouette scores were used to assist with clustering. Clustering was performed with leiden with resolutions 1.0 for yAL and 0.9 for yDR. Cells were labeled according to marker genes.

IGF target gene UCell scores (Andreatta & Carmona, 2026) were calculated using the HALLMARK_MTORC1_SIGNALING gene set from MSigDB (D. G. Howe et al., 2018) as a proxy for downstream IGF pathway activity (hereafter termed mTORC1 UCell score). For pseudotime analysis, clusters corresponding to erythroid progenitors (ErP), B cells, and dendritic cells were removed from both yAL and yDR datasets prior to trajectory inference. Diffusion pseudotime (DPT) (Haghverdi et al., 2016) was then calculated using Scanpy. The root cell was defined based on expression of the hematopoietic stem cell markers *Mecom* and *Mpl*. For yDR, the root cell corresponded to the cell with the highest combined expression of *Mecom* and *Mpl*. For yAL, the root cell was defined as the cell with the fourth highest combined expression because the three highest-ranking cells were either outside the HSC cluster or located at the boundary between the HSC cluster and neighboring populations.

Cells labeled as HSC, MPP, and MPP S-phase (hereafter termed MPP Cycling) were isolated for pseudotime analysis. Cells were ordered according to DPT and aggregated into 80 pseudocells as previously described in (Rommelfanger et al., 2025). Pseudocell expression was defined as the average expression of all cells assigned to each pseudocell bin. Pseudocell-level expression profiles were then computed for *Mecom*, *Cdk6*, and the mTORC1 UCell score across pseudotime for each experimental condition.

## Acknowledgements

We would like to thank members of the MacLean and Rudolph labs for helpful discussions. A.L.M. acknowledges support from the National Science Foundation (GR1066485) and the National Institutes of Health (R35GM143019).

## Data availability statement

All of the code and analysis scripts associated with this manuscript, developed in MATLAB (Natick, MA) and Python, are available on GitHub at https://github.com/maclean-lab/mecom-toggle.

## Author Contributions

**Conceptualization**: JAM, KLR, ALM. **Methodology**: JAM, AD, ALM. **Software**: JAM, AD, MM. **Investigation**: JAM, AD, MM, ALM. **Formal analysis**: JAM, AD, ALM. **Writing – original draft**: JAM, AD, MM, ALM. **Writing – reviewing and editing**: JAM, AD, MM, KLR, ALM. **Funding acquisition**: KLR, ALM. **Supervision**: KLR, ALM.

## References

1. Andreatta, M., & Carmona, S. J. (2026). UCell and pyUCell: Single-cell gene signature scoring for R and python. Bioinformatics, 42(2).

2. Benveniste, P., Frelin, C., Janmohamed, S., Barbara, M., Herrington, R., Hyam, D., & Iscove, N. N. (2010). Intermediate-term hematopoietic stem cells with extended but time-limited reconstitution potential. Cell Stem Cell, 6(1), 48–58.

3. Braal, C. L., Jongbloed, E. M., Wilting, S. M., Mathijssen, R. H. J., Koolen, S. L. W., & Jager, A. (2021). Inhibiting CDK4/6 in breast cancer with palbociclib, ribociclib, and abemaciclib: Similarities and differences. Drugs, 81(3), 317–331.

4. Calvanese, V., Capellera-Garcia, S., Ma, F., Fares, I., Liebscher, S., Ng, E. S., Ekstrand, S., Aguadé-Gorgorió, J., Vavilina, A., Lefaudeux, D., Nadel, B., Li, J. Y., Wang, Y., Lee, L. K., Ardehali, R., Iruela-Arispe, M. L., Pellegrini, M., Stanley, E. G., Elefanty, A. G., … Mikkola, H. K. A. (2022). Mapping human haematopoietic stem cells from haemogenic endothelium to birth. Nature, 604(7906), 534–540.

5. Cheng, T., Rodrigues, N., Shen, H., Yang, Y., Dombkowski, D., Sykes, M., & Scadden, D. T. (2000). Hematopoietic stem cell quiescence maintained by p21cip1/waf1. Science, 287(5459), 1804–1808.

6. Cislo, D. J., Delás, M. J., Briscoe, J., & Siggia, E. D. (2025). Reconstructing waddington’s landscape from data. Proc. Natl. Acad. Sci. U. S. A., 122(49), e2521762122.

7. de Haan, G., & Lazare, S. S. (2018). Aging of hematopoietic stem cells. Blood, 131(5), 479–487.

8. Dey, A., & MacLean, A. L. (2025). Transition paths across the epithelial-mesenchymal transition landscape are dictated by network logic. Development, 152(11), dev204583.

9. Duddu, A. S., Jolly, M. K., & Raju, A. (2025). Distinct geometrical landscapes distinguish between modes of tristability in gene regulatory networks.

10. Ediriwickrema, A., Nakauchi, Y., Fan, A. C., Köhnke, T., Hu, X., Luca, B. A., Kim, Y., Ramakrishnan, S., Nakamoto, M., Karigane, D., Linde, M. H., Azizi, A., Newman, A. M., Gentles, A. J., & Majeti, R. (2025). A single-cell framework identifies functionally and molecularly distinct multipotent progenitors in adult human hematopoiesis. Cell Rep., 44(9), 116236.

12. Fujino, T., Asada, S., Goyama, S., & Kitamura, T. (2022). Mechanisms involved in hematopoietic stem cell aging. Cell. Mol. Life Sci., 79(9), 473.

13. Grockowiak, E., Korn, C., Rak, J., Lysenko, V., Hallou, A., Panvini, F. M., Williams, M., Fielding, C., Fang, Z., Khatib-Massalha, E., Garcıa-Garcıa, A., Li, J., Khorshed, R. A., González-Antón, S., Baxter, E. J., Kusumbe, A., Wilkins, B. S., Green, A., Simons, B. D., … Méndez-Ferrer, S. (2023). Different niches for stem cells carrying the same oncogenic driver affect pathogenesis and therapy response in myeloproliferative neoplasms. *Nat*. Cancer, 4(8), 1193–1209.

14. Haghverdi, L., Büttner, M., Wolf, F. A., Buettner, F., & Theis, F. J. (2016). Diffusion pseudotime robustly reconstructs lineage branching. Nat. Methods, 13(10), 845–848.

15. He, S., Roberts, P. J., Sorrentino, J. A., Bisi, J. E., Storrie-White, H., Tiessen, R. G., Makhuli, K. M., Wargin, W. A., Tadema, H., van Hoogdalem, E.-J., Strum, J. C., Malik, R., & Sharpless, N. E. (2017). Transient CDK4/6 inhibition protects hematopoietic stem cells from chemotherapy-induced exhaustion. Sci. Transl. Med., 9(387), eaal3986.

16. Howe, A., & Mani, M. (2025). Learning geometric models for developmental dynamics. Phys. Rev. X., 15(3).

17. Howe, D. G., Blake, J. A., Bradford, Y. M., Bult, C. J., Calvi, B. R., Engel, S. R., Kadin, J. A., Kaufman, T. C., Kishore, R., Laulederkind, S. J. F., Lewis, S. E., Moxon, S. A. T., Richardson, J. E., & Smith, C. (2018). Model organism data evolving in support of translational medicine. Lab Anim. (NY), 47(10), 277–289.

18. Huang, S., Guo, Y.-P., May, G., & Enver, T. (2007). Bifurcation dynamics in lineage-commitment in bipotent progenitor cells. Dev. Biol., 305(2), 695–713.

19. Jakobsen, N. A., Turkalj, S., Zeng, A. G. X., Stoilova, B., Metzner, M., Rahmig, S., Nagree, M. S., Shah, S., Moore, R., Usukhbayar, B., Angulo Salazar, M., Gafencu, G.-A., Kennedy, A., Newman, S., Kendrick, B. J. L., Taylor, A. H., Afinowi-Luitz, R., Gundle, R., Watkins, B., … Vyas, P. (2024). Selective advantage of mutant stem cells in human clonal hematopoiesis is associated with attenuated response to inflammation and aging. Cell Stem Cell, 31(8), 1127–1144.e17.

20. Johnson, S. M., Torrice, C. D., Bell, J. F., Monahan, K. B., Jiang, Q., Wang, Y., Ramsey, M. R., Jin, J., Wong, K.-K., Su, L., Zhou, D., & Sharpless, N. E. (2010). Mitigation of hematologic radiation toxicity in mice through pharmacological quiescence induced by CDK4/6 inhibition. J. Clin. Invest., 120(7), 2528–2536.

21. Kollmann, K., Heller, G., Schneckenleithner, C., Warsch, W., Scheicher, R., Ott, R. G., Schäfer, M., Fajmann, S., Schlederer, M., Schiefer, A.-I., Reichart, U., Mayerhofer, M., Hoeller, C., Zöchbauer-Müller, S., Kerjaschki, D., Bock, C., Kenner, L., Hoefler, G., Freissmuth, M., … Sexl, V. (2013). A kinase-independent function of CDK6 links the cell cycle to tumor angiogenesis. Cancer Cell, 24(2), 167–181.

22. Kramer, A., & Challen, G. A. (2017). The epigenetic basis of hematopoietic stem cell aging. Semin. Hematol., 54(1), 19–24.

23. Kreger, J., Mooney, J. A., Shibata, D., & MacLean, A. L. (2024). Developmental hematopoietic stem cell variation explains clonal hematopoiesis later in life. Nat. Commun., 15(1), 10268.

24. Kuznetsov, Y. A. (2023). Elements of applied bifurcation theory. Springer International Publishing.

25. Laurenti, E., Frelin, C., Xie, S., Ferrari, R., Dunant, C. F., Zandi, S., Neumann, A., Plumb, I., Doulatov, S., Chen, J., April, C., Fan, J.-B., Iscove, N., & Dick, J. E. (2015). CDK6 levels regulate quiescence exit in human hematopoietic stem cells. Cell Stem Cell, 16(3), 302–313.

26. Liberzon, A., Birger, C., Thorvaldsdóttir, H., Ghandi, M., Mesirov, J. P., & Tamayo, P. (2015). The molecular signatures database (MSigDB) hallmark gene set collection. Cell Syst., 1(6), 417–425.

27. MacLean, A. L., & Rudolph, K. L. (2024). Hematopoietic stem cell aging by the niche. Blood, 144(4), 347–348.

28. MacLean, A. L., Smith, M. A., Liepe, J., Sim, A., Khorshed, R., Rashidi, N. M., Scherf, N., Krinner, A., Roeder, I., Lo Celso, C., & Stumpf, M. P. H. (2017). Single cell phenotyping reveals heterogeneity among hematopoietic stem cells following infection. Stem Cells, 35(11), 2292–2304.

29. MacLean, A. L., Hong, T., & Nie, Q. (2018). Exploring intermediate cell states through the lens of single cells. Curr. Opin. Syst. Biol., 9, 32–41.

30. Maurer, B., Brandstoetter, T., Kollmann, S., Sexl, V., & Prchal-Murphy, M. (2021). Inducible deletion of CDK4 and CDK6 – deciphering CDK4/6 inhibitor effects in the hematopoietic system. Haematologica, 106(10), 2624–2632.

31. Mayer, I. M., Doma, E., Klampfl, T., Prchal-Murphy, M., Kollmann, S., Schirripa, A., Scheiblecker, L., Zojer, M., Kunowska, N., Gebrail, L., Shaw, L. E., Mann, U., Farr, A., Grausenburger, R., Heller, G., Zebedin-Brandl, E., Farlik, M., Malumbres, M., Sexl, V., & Kollmann, K. (2024). Kinase-inactivated CDK6 preserves the long-term functionality of adult hematopoietic stem cells. Blood, 144(2), 156–170.

32. McInnes, L., Healy, J., & Melville, J. (2018). UMAP: Uniform manifold approximation and projection for dimension reduction.

33. Mejia-Ramirez, E., & Florian, M. C. (2020). Understanding intrinsic hematopoietic stem cell aging. Haematologica, 105(1), 22–37.

34. Mendelson, A., & Frenette, P. S. (2014). Hematopoietic stem cell niche maintenance during homeostasis and regeneration. Nat. Med., 20(8), 833–846.

35. Moris, N., Pina, C., & Arias, A. M. (2016). Transition states and cell fate decisions in epigenetic landscapes. Nat. Rev. Genet., 17(11), 693–703.

36. Morrison, L., Loibl, S., & Turner, N. C. (2024). The CDK4/6 inhibitor revolution – a game-changing era for breast cancer treatment. Nat. Rev. Clin. Oncol., 21(2), 89–105.

37. Morrison, S. J., & Scadden, D. T. (2014). The bone marrow niche for haematopoietic stem cells. Nature, 505(7483), 327–334.

38. Nakamura-Ishizu, A., Takizawa, H., & Suda, T. (2014). The analysis, roles and regulation of quiescence in hematopoietic stem cells. Development, 141(24), 4656–4666.

39. Raju, A., & Siggia, E. D. (2023). A geometrical perspective on development. Dev. Growth Differ., 65(5), 245–254.

40. Raju, A., & Siggia, E. D. (2024). A geometrical model of cell fate specification in the mouse blastocyst. Development, 151(8), dev202467.

41. Rand, D. A., Raju, A., Sáez, M., Corson, F., & Siggia, E. D. (2021). Geometry of gene regulatory dynamics. Proc. Natl. Acad. Sci. U. S. A., 118(38), e2109729118.

42. Rommelfanger, M. K., & MacLean, A. L. (2021). A single-cell resolved cell-cell communication model explains lineage commitment in hematopoiesis. Development, 148(24), dev199779.

43. Rommelfanger, M. K., Behrends, M., Chen, Y., Martinez, J., Kurella, N., Geisler, N., Guturu, D., Bens, M., Xiong, L., Xiang, Z., Rudolph, K. L., & MacLean, A. L. (2025). Gene regulatory network inference with popinfer reveals dynamic regulation of hematopoietic stem cell quiescence. iScience, (114010), 114010.

44. Sáez, M., Briscoe, J., & Rand, D. A. (2022). Dynamical landscapes of cell fate decisions. Interface Focus, 12(4), 20220002.

45. Safina, K., & van Galen, P. (2024). New frameworks for hematopoiesis derived from single-cell genomics. Blood, 144(10), 1039–1047.

46. Satija, R., Farrell, J. A., Gennert, D., Schier, A. F., & Regev, A. (2015). Spatial reconstruction of single-cell gene expression data. Nat. Biotechnol., 33(5), 495–502.

47. Scheicher, R., Hoelbl-Kovacic, A., Bellutti, F., Tigan, A.-S., Prchal-Murphy, M., Heller, G., Schneckenleithner, C., Salazar-Roa, M., Zöchbauer-Müller, S., Zuber, J., Malumbres, M., Kollmann, K., & Sexl, V. (2015). CDK6 as a key regulator of hematopoietic and leukemic stem cell activation. Blood, 125(1), 90–101.

48. Sparta, B., Hamilton, T., Hughes, S., Natesan, G., & Deeds, E. (2022). A lack of distinct cell identities in single-cell measurements: Revisiting waddington’s landscape.

49. Suo, M., Rommelfanger, M. K., Chen, Y., Amro, E. M., Han, B., Chen, Z., Szafranski, K., Chakkarappan, S. R., Boehm, B. O., MacLean, A. L., & Rudolph, K. L. (2022). Age-dependent effects of igf2bp2 on gene regulation, function, and aging of hematopoietic stem cells in mice. Blood, 139(17), 2653–2665.

50. Thom, R. (1975). Structural stability and morphogenesis: An outline of a general theory of models.

51. Tirosh, I., Izar, B., Prakadan, S. M., Wadsworth, M. H., 2nd, Treacy, D., Trombetta, J. J., Rotem, A., Rodman, C., Lian, C., Murphy, G., Fallahi-Sichani, M., Dutton-Regester, K., Lin, J.-R., Cohen, O., Shah, P., Lu, D., Genshaft, A. S., Hughes, T. K., Ziegler, C. G. K., … Garraway, L. A. (2016). Dissecting the multicellular ecosystem of metastatic melanoma by single-cell RNA-seq. Science, 352(6282), 189–196.

52. Toghani, D., Gupte, S., Zeng, S., Mahammadov, E., Crosse, E. I., Seyedhassantehrani, N., Burns, C., Gravano, D., Radtke, S., Kiem, H.-P., Rodriguez, S., Carlesso, N., Pradeep, A., Georgiades, A., Lucas, F., Wilson, N. K., Kinston, S. J., Göttgens, B., Zong, L., … Silberstein, L. (2025). Niche-derived semaphorin 4A safeguards functional identity of myeloid-biased hematopoietic stem cells. *Nat*. Aging, 5(4), 558–575.

53. Tümpel, S., & Rudolph, K. L. (2019). Quiescence: Good and bad of stem cell aging. Trends Cell Biol., 29(8), 672–685.

54. Voit, R. A., Tao, L., Yu, F., Cato, L. D., Cohen, B., Fleming, T. J., Antoszewski, M., Liao, X., Fiorini, C., Nandakumar, S. K., Wahlster, L., Teichert, K., Regev, A., & Sankaran, V. G. (2023). A genetic disorder reveals a hematopoietic stem cell regulatory network co-opted in leukemia. Nat. Immunol., 24(1), 69–83.

55. Wang, J. C. Y., & Dick, J. E. (2005). Cancer stem cells: Lessons from leukemia. Trends Cell Biol., 15(9), 494–501.

56. Weng, C., Yu, F., Yang, D., Poeschla, M., Liggett, L. A., Jones, M. G., Qiu, X., Wahlster, L., Caulier, A., Hussmann, J. A., Schnell, A., Yost, K. E., Koblan, L. W., Martin-Rufino, J. D., Min, J., Hammond, A., Ssozi, D., Bueno, R., Mallidi, H., … Sankaran, V. G. (2024). Deciphering cell states and genealogies of human haematopoiesis. Nature, 627(8003), 389–398.

57. Wilson, A., Laurenti, E., Oser, G., van der Wath, R. C., Blanco-Bose, W., Jaworski, M., Offner, S., Dunant, C. F., Eshkind, L., Bockamp, E., Lió, P., Macdonald, H. R., & Trumpp, A. (2008). Hematopoietic stem cells reversibly switch from dormancy to self-renewal during homeostasis and repair. Cell, 135(6), 1118–1129.

58. Wolf, F. A., Angerer, P., & Theis, F. J. (2018). SCANPY: Large-scale single-cell gene expression data analysis. Genome Biol., 19(1), 15.

59. Wolock, S. L., Lopez, R., & Klein, A. M. (2019). Scrublet: Computational identification of cell doublets in single-cell transcriptomic data. Cell Syst., 8(4), 281–291.e9.

60. Yampolskaya, M., Ikonomou, L., & Mehta, P. (2025). Finding signatures of low-dimensional geometric landscapes in high-dimensional cell fate transitions, bioRxiv.

61. Yang, L., Bryder, D., Adolfsson, J., Nygren, J., Månsson, R., Sigvardsson, M., & Jacobsen, S. E. W. (2005). Identification of Lin(-)Sca1(+)kit(+)CD34(+)Flt3-short-term hematopoietic stem cells capable of rapidly reconstituting and rescuing myeloablated transplant recipients. Blood, 105(7), 2717–2723.

62. Yokomizo, T., Ideue, T., Morino-Koga, S., Tham, C. Y., Sato, T., Takeda, N., Kubota, Y., Kurokawa, M., Komatsu, N., Ogawa, M., Araki, K., Osato, M., & Suda, T. (2022). Independent origins of fetal liver haematopoietic stem and progenitor cells. Nature, 609(7928), 779–784.

63. Young, K., Eudy, E., Bell, R., Loberg, M. A., Stearns, T., Sharma, D., Velten, L., Haas, S., Filippi, M.-D., & Trowbridge, J. J. (2021). Decline in IGF1 in the bone marrow microenvironment initiates hematopoietic stem cell aging. Cell Stem Cell, 28(8), 1473–1482.e7.

64. Young, K. A., Telpoukhovskaia, M. A., Hofmann, J., Mistry, J. J., Kokkaliaris, K. D., & Trowbridge, J. J. (2024). Variation in mesenchymal KITL/SCF and IGF1 expression in middle age underlies steady-state hematopoietic stem cell aging. Blood, 144(4), 378–391.

65. Zeng, A. G. X., Nagree, M. S., Jakobsen, N. A., Shah, S., Varesi, A., Kang, J. R. W., Murison, A., Cheong, J.-G., Turkalj, S., Zhang, X., Radtke, F. A., Abera, T.-A., Lim, I. N. X., Jin, L., Araújo, J., Aguilar-Navarro, A. G., Parris, D., McLeod, J., Kim, H., … Xie, S. Z. (2026). Human haematopoietic stem cells remember inflammatory stress. Nature.

