## Supplementary Tables and Figures for "A *Mecom—Cdk6* toggle switch governs hematopoietic stem-to-multipotent fate transitions via distinct multistable landscapes"

Supplemental Table 1: *Additional parameter sets exhibiting tetrastability.* Two additional parameterizations of the MECOM–CDK6 model that produce tetrastability in the baseline model. These parameter sets were used to assess whether the transition geometries observed, following weakening of MECOM self-activation, are robust to parameter selection. Corresponding bifurcation diagrams and minimum action paths are shown in Supplemental Fig. 4.

| Parameter | Parameter Set 2 | Parameter Set 3 |
|---|---|---|
| $k_X = k_Y$ | 10.68 | 14.50 |
| $\beta_X = \beta_Y$ | 465.18 | 223.42 |
| $\delta_X = \delta_Y$ | 0.011 | 0.020 |
| $\gamma_X = \gamma_Y$ | 232.43 | 73.93 |
| $k_{X0} = k_{Y0}$ | 0.345 | 0.588 |
| $n_1 = n_4$ | 1 | 4 |
| $n_2 = n_5$ | 6 | 3 |
| $n_3 = n_6$ | 8 | 9 |
| $\alpha_X = \alpha_Y$ | 60.30 | 53.05 |

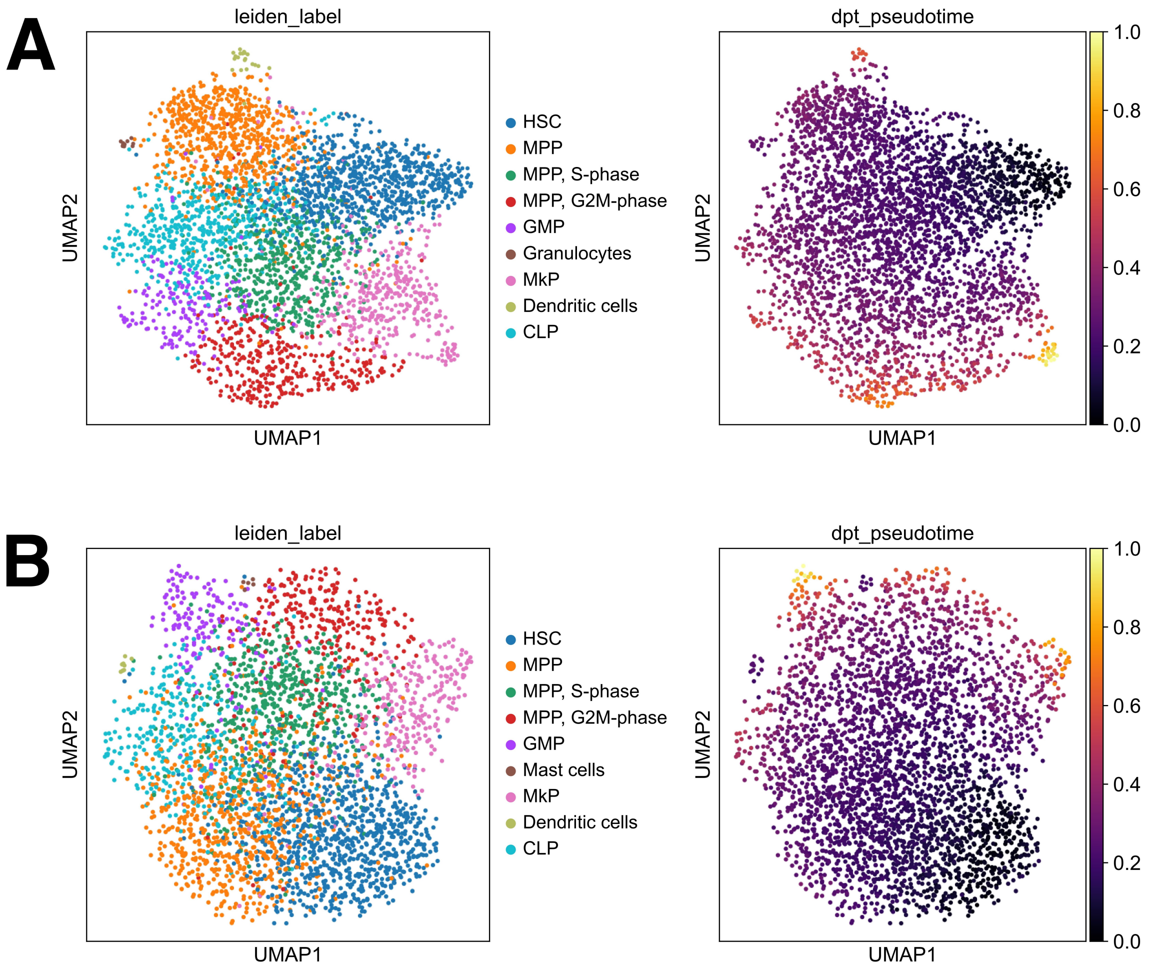


Supplemental Figure 1: *Annotated cell types and diffusion pseudotime projected onto UMAPs of young hematopoietic cells.* **(A)** UMAP colored by cell type (left) and DPT pseudotime (right) for young ad libitum samples. **(B)** Same as (A), but for young diet-restricted samples.

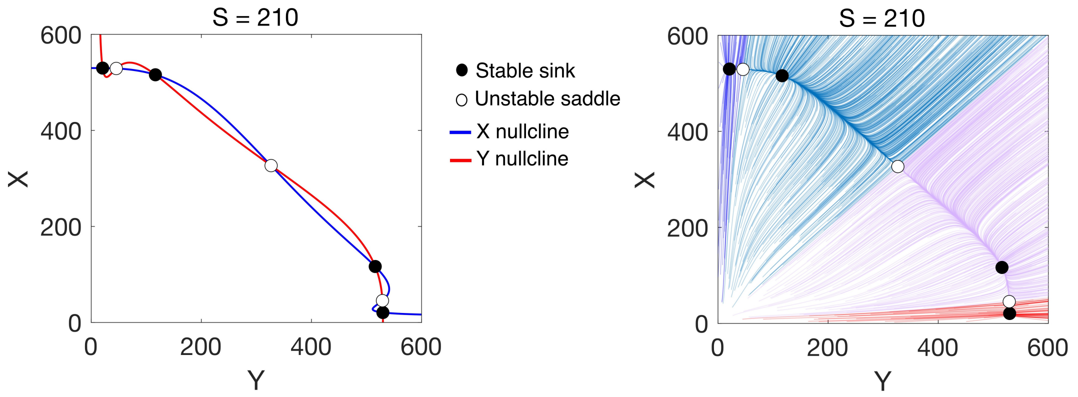


Supplemental Figure 2: *The baseline tetrastable regime contains extensive intermediate-state basins separating stem and multipotent attractors.* Nullclines (left) and phase-plane trajectories (right) for the baseline model at $S = 210$. Blue trajectories converge to the MECOM-high (stem) state, red trajectories converge to the CDK6-high (multipotent) state, and light blue and purple trajectories converge to the two intermediate states.

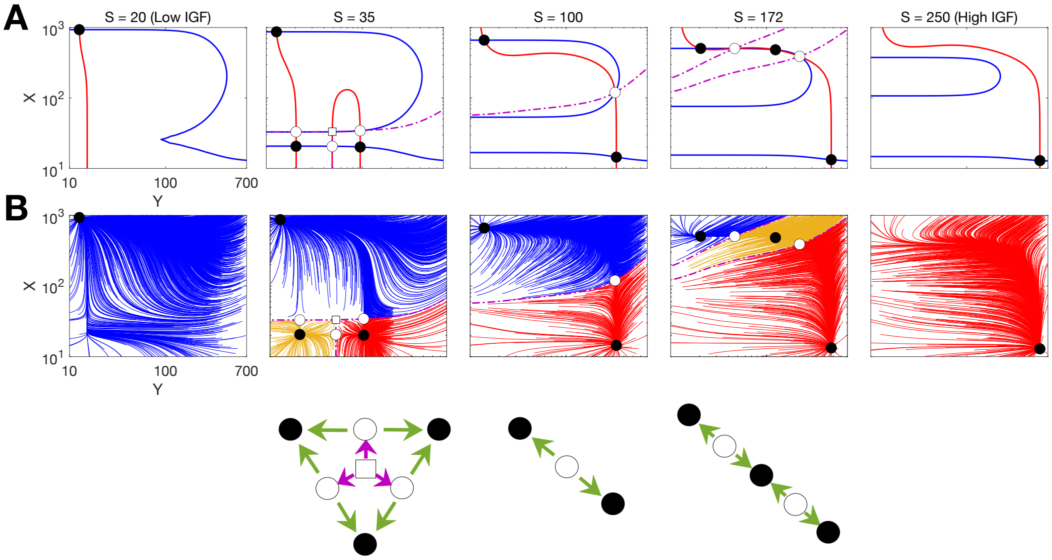


Supplemental Figure 3: *Evolution of landscape geometry reveals how increasing IGF reshapes transitions following weaked Mecom self-activation.* **(A)** Phase-plane nullclines at representative IGF values ($S$ = 20-250) for 2.5-fold decrease in $\alpha_x$. *Cdk6* nullclines are shown in red and *Mecom* nullclines in blue. Filled black circles denote stable steady states, open circles denote saddle points, open squares denote source points, and purple dashed curves denote separatrices. **(B)** Phase-plane trajectory flows for representative IGF values in (A). Red trajectories converge to the *Cdk6*-high (multipotent) state, blue trajectories converge to the *Mecom*-high (stem) state, and yellow trajectories converge to intermediate states. Schematic geometries corresponding to selected landscapes are shown below. Green arrows denote unstable manifolds and magenta arrows denote stable manifolds.

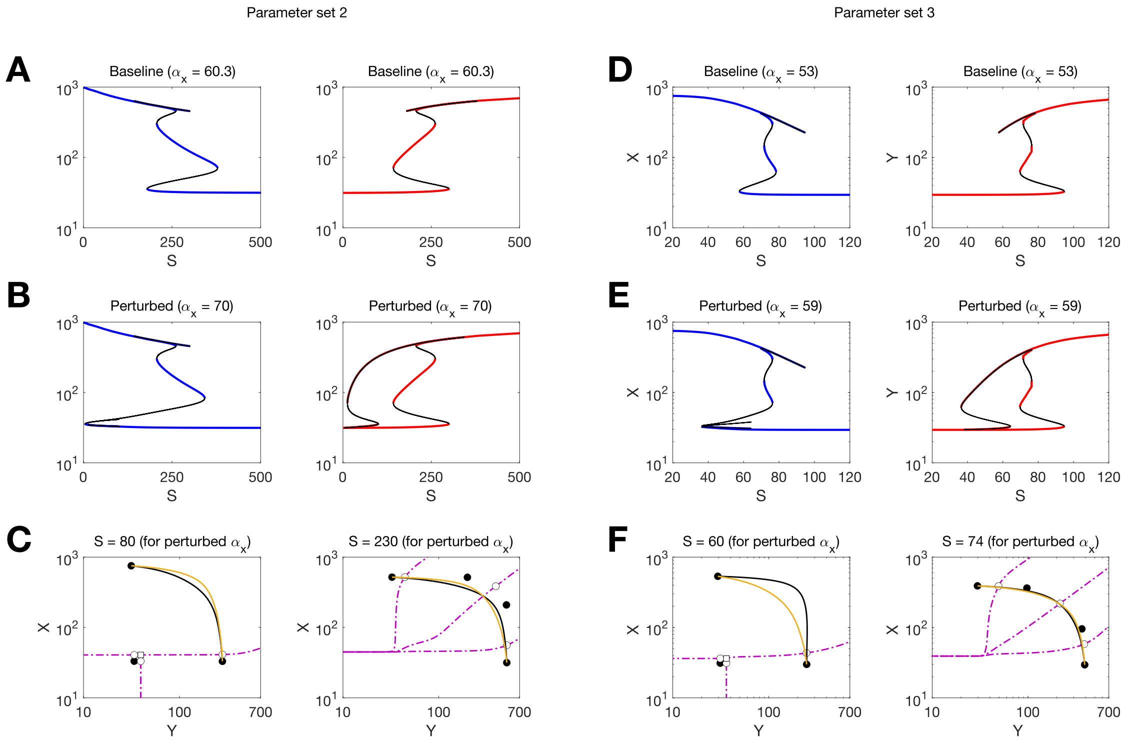


Supplemental Figure 4: *Weakening MECOM self-activation produces similar transition geometries across additional tetrastable parameter sets.* **(A)** Baseline one-parameter bifurcation diagrams for Parameter Set 2 (Supplemental Table 1) showing MECOM ($X$, blue) and CDK6 ($Y$, red) steady states as a function of IGF signaling ($S$). **(B)** Same as (A) following weakening of MECOM self-activation by increasing $\alpha_x$. As in the primary parameter set, the tetrastable region separates into distinct low- and high-IGF tristable regimes. **(C)** Minimum action paths for the perturbed Parameter Set 2 at representative low-IGF (left) and high-IGF (right) conditions. **(D)** Same as (A), but for Parameter Set 3. **(E)** Same as (B), but for Parameter Set 3. **(F)** Same as (C), but for Parameter Set 3.

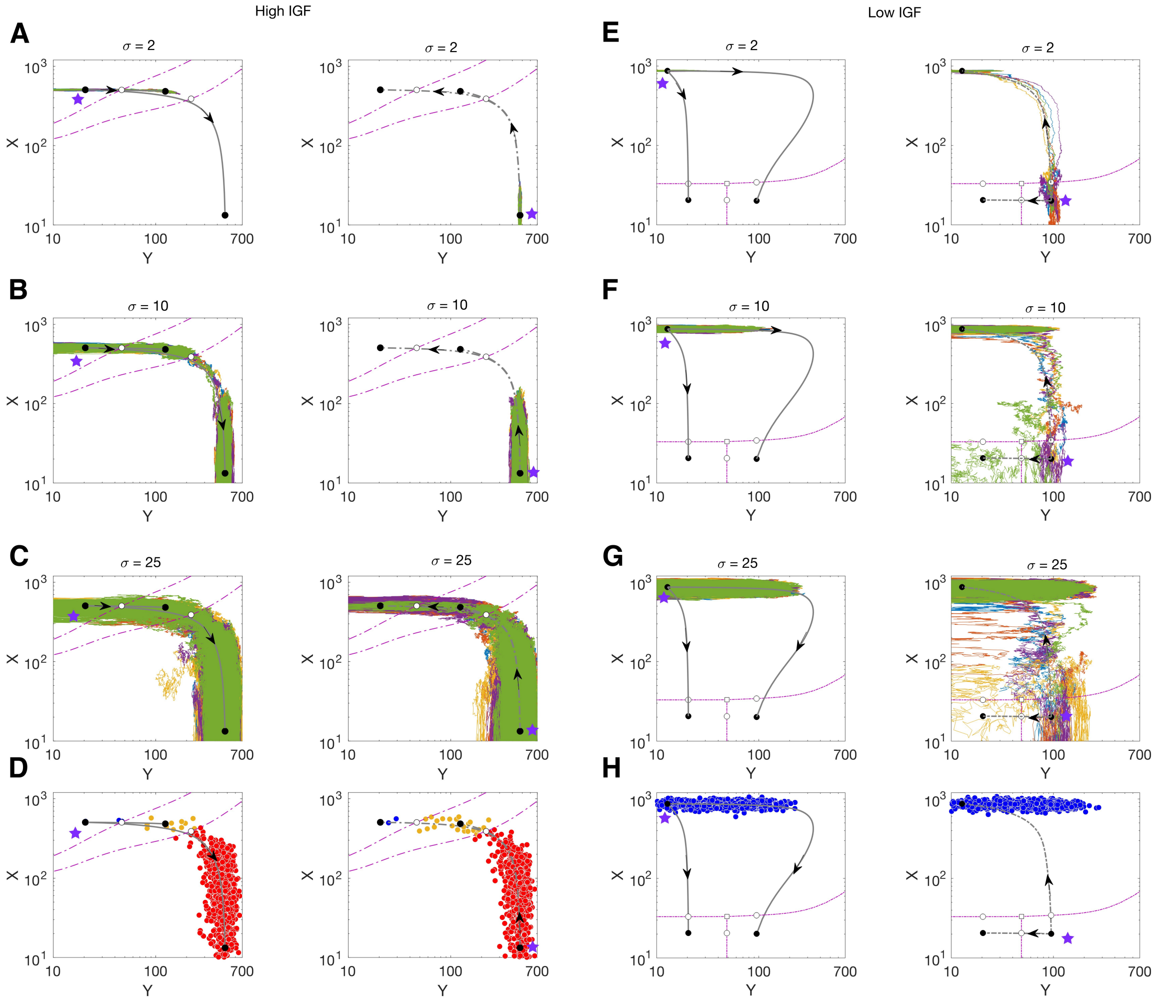


Supplemental Figure 5: *Stochastic trajectories and realizations reveal that intermediate states mediate transitions at high IGF but are not required at low IGF.* **(A)** High-IGF regime for a 2.5-fold increase in MECOM self-activation ($\alpha_x = 217.5$). Separatrices (purple dashed curves) and minimum action paths (black curves) are overlaid with five stochastic trajectories (each represented by a different color) initialized near the *Mecom*-high (left) or *Cdk6*-high (right) attractor (purple star) for noise strength $\sigma = 2$. **(B)** Same as (A), but for $\sigma = 10$. **(C)** Same as (A), but for $\sigma = 25$. **(D)** Cell states at the final simulation time (10,000 stochastic trajectories at time $t = 10^4$) under the conditions shown in (C). Blue points indicate trajectories ending in the *Mecom*-high state, yellow points indicate trajectories ending in an intermediate state, and red points indicate trajectories ending in the *Cdk6*-high state. **(E)** Same as (A), but for the low-IGF regime. **(F)** Same as (E), but for $\sigma = 10$. **(G)** Same as (E), but for $\sigma = 25$. **(H)** Same as (D), but for the conditions shown in (G).

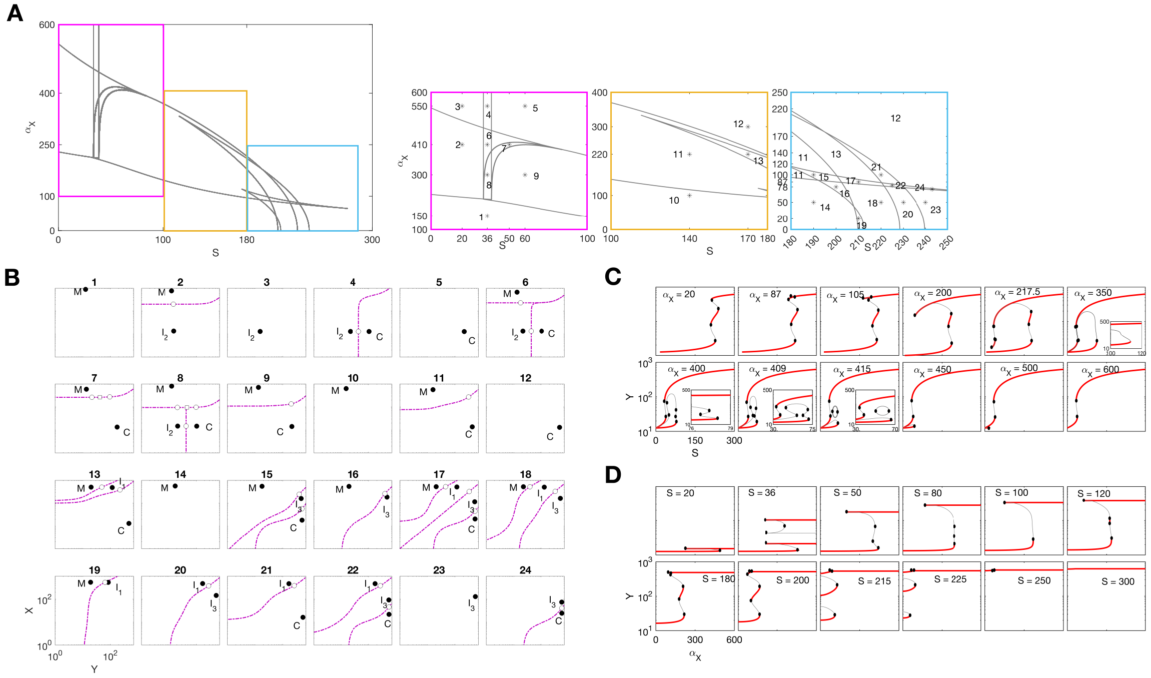


Supplemental Figure 6: *Two-parameter bifurcation analysis identifies distinct landscape organizations across IGF signaling and Mecom self-activation levels.* **(A)** Two-parameter bifurcation diagram in the $(S, \alpha_x)$ plane, where $S$ denotes IGF pathway signaling and $\alpha_x$ controls *Mecom* self-activation. Colored boxes highlight representative low-IGF (purple), intermediate-IGF (yellow), and high-IGF (blue) regions. Numbers indicate distinct regions of parameter space associated with different landscape organizations. **(B)** Attractors and basin boundaries corresponding to the numbered regions in (A). Filled black circles denote stable steady states, open circles denote saddle points, open squares denote source points, and purple dashed curves denote separatrices. $M$ indicates the *Mecom*-high (stem) state, $C$ indicates the *Cdk6*-high (multipotent) state, and $I_1$, $I_2$, and $I_3$ denote intermediate states. **(C)** One-parameter bifurcation diagrams of *Cdk6* obtained by varying $S$ while holding $\alpha_x$ fixed at the indicated values. Red curves denote stable steady states and black curves denote unstable steady states. Numbered labels correspond to the regions identified in (A). **(D)** Same as (C), but varying $\alpha_x$ while holding $S$ fixed at the indicated values.

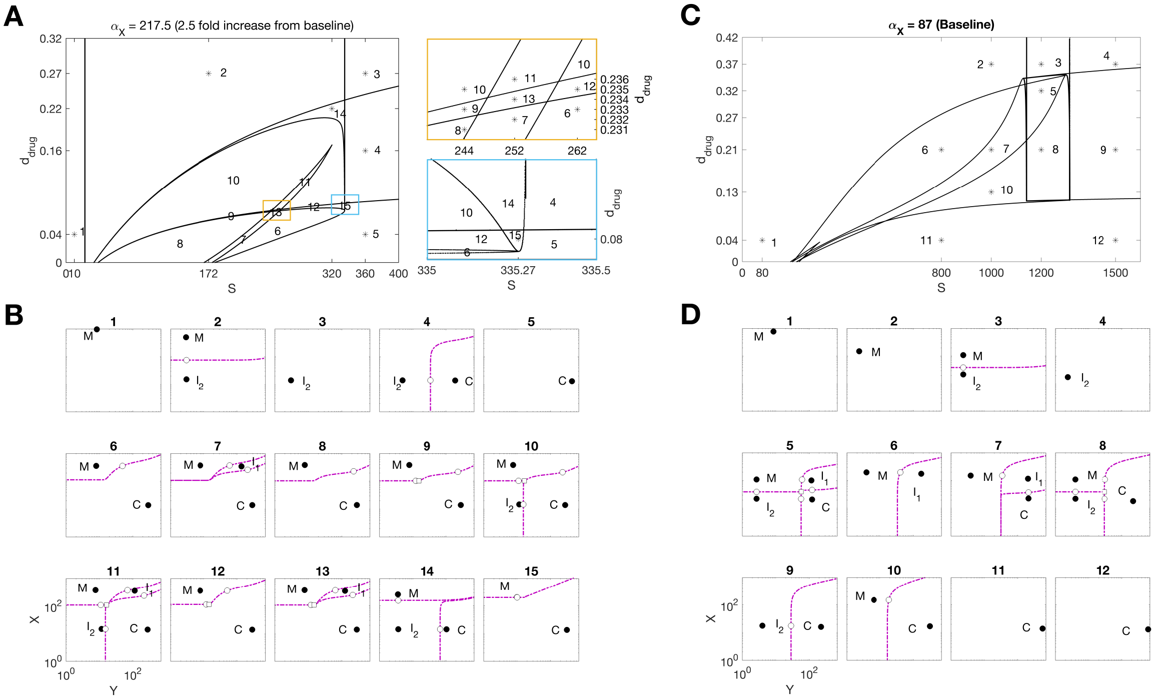

Supplemental Figure 7: *Cdk6 inhibition reorganizes landscape geometries across IGF signaling and Mecom self-activation levels.* **(A)** Two-parameter bifurcation diagram in the $(S, d_{drug})$ plane for a 2.5-fold increase in $\alpha_x$ relative to baseline ($\alpha_x = 217.5$), where $S$ denotes IGF pathway signaling and $d_{drug}$ represents drug-induced degradation of *Cdk6*. Numbers indicate distinct regions of parameter space associated with different landscape organizations. **(B)** Attractors and basin boundaries corresponding to the numbered regions in (A). Filled black circles denote stable steady states, open circles denote saddle points, open squares denote source points, and purple dashed curves denote separatrices. $M$ indicates the *Mecom*-high (stem) state, $C$ indicates the *Cdk6*-high (multipotent) state, and $I_1$ and $I_2$ denote intermediate states. **(C)** Same as (A), but for the baseline value of *Mecom* self-activation ($\alpha_x = 87$). **(D)** Same as (B), but corresponding to the numbered regions identified in (C).
